# Reduced γ-glutamyl hydrolase activity likely contributes to high folate levels in Periyakulam-1 tomato

**DOI:** 10.1101/2022.07.30.502140

**Authors:** Kamal Tyagi, Anusha Sunkum, Prateek Gupta, Himabindu Vasuki Kilambi, Yellamaraju Sreelakshmi, Rameshwar Sharma

## Abstract

Tomato cultivars show wide variation in nutraceutical folate in ripe fruits, yet the loci regulating folate levels in fruits remain unexplored. To decipher regulatory points, we compared two contrasting tomato cultivars: Periyakulam-1 (PKM-1) with high folate and Arka Vikas (AV) with low folate. The progression of ripening in PKM-1 was nearly similar to AV but had substantially lower ethylene emission. In parallel, the levels of phytohormones salicylic acid, ABA, and jasmonic acid were substantially lower than AV. The fruits of PKM-1 were also metabolically distinct from AV, with upregulation of several amino acids. Consistent with higher °Brix levels, the red ripe fruits also showed upregulation of sugars and sugar-derived metabolites. In parallel with higher folate, PKM-1 fruits also had higher carotenoid levels, especially lycopene and β-carotene. Transcript levels of genes encoding folate biosynthesis did not show a perceptible difference in relative expression. The proteome analysis showed upregulation of carotenoid sequestration and folate metabolism-related proteins in PKM-1. The deglutamylation pathway mediated by γ-glutamyl hydrolase (GGH) was substantially reduced in PKM-1 at the red-ripe stage. The red-ripe fruits had reduced transcript levels of GGHs and lower GGH activity than AV. Conversely, the percent polyglutamylation of folate was much higher in PKM-1. Our analysis indicates the regulation of GGH activity as a potential target to elevate folate levels in tomato fruits.

## Introduction

Tomato is the sixth most important globally grown crop. Tomato is also a nutritionally important crop as it is a rich source of several nutraceuticals, particularly antioxidant lycopene and β-carotene, a precursor of vitamin A (Chaudhary et al., 2018). Though tomato leaves are enriched in vitamin B9, commonly called folate, the fruits have a moderate folate level (Tyagi et al., 2015). Folate is an essential vitamin for all organisms; however, animals cannot synthesize folate and obtain it from plant-derived food. The staple cereal crops, wheat and rice have low folate levels, necessitating the fortification of foods with folic acid to meet the RDA (recommended dietary allowance) (Blancquaert et al., 2014; Jiang et al., 2021). To obviate the folate fortification, efforts are directed to biofortify the important crops with folate (Gorelova et al., 2017a).

The strategy to biofortify the folate in the edible portion of the crop plants has focused on two approaches, germplasm diversity and metabolic engineering. The studies on transgenic manipulation of folate biosynthesis in tomato revealed that upregulation of both GTP cyclohydrolase I (GCHI), and aminodeoxychorismate synthase (ADCS), the key enzymes synthesizing pterin and p-aminobenzoic acid (pABA), respectively, was essential for folate biofortification (de la Garza et al., 2007). Alike tomato, the transgenic upregulation of GCHI and ADCS in rice also led to folate enrichment (Storozhenko et al., 2007). Contrastingly, in potato tubers, the overexpression of GCHI and ADCS did not improve folate levels, highlighting species-specific regulation of folate biosynthesis (Blancquaert et al., 2013). The inclusion of two additional genes pyrophosphokinase/dihydropteroate synthase (HPPK/DHPS) and folylpolyglutamate synthetase (FPGS), enabled a 12-fold enrichment of folate in potato (De Lepeleire et al., 2018). The protection of folate polyglutamylation by lowering γ-glutamyl hydrolase (GGH) level moderately improved the folate levels in tomato fruits (Akhtar et al., 2010).

The analysis of folate levels in different crop species revealed a limited range of variation across the germplasm. An analysis of folate levels in 175 wheat genotypes revealed 2.12- and 2.27-fold variation in winter wheat and spring wheat, respectively (Piironen et al., 2008). An analysis of 67 accessions in spinach showed a 3.2-fold variation in folate levels (Shohag et al., 2011). In 125 accessions of tomato, the folate level in red-ripe fruits varied from 13.8-45.8 μg/100 g FW (Fresh weight), while in mature-green fruits, it ranged from 12.5-70.9 μg/100 g FW. In most accessions, folate levels declined during fruit ripening (Upadhyaya et al., 2017). The decline in folate level may be causally associated with reduced expression of GCHI and ADCS with the progression of ripening in tomato fruits (Waller et al., 2010).

The limited diversity in folate levels in germplasm may be associated with the absence of polymorphism in genes regulating folate biosynthesis. In tomato, analysis of 391 accessions revealed the paucity of SNPs in exons of *GCHI*, *ADCS*, and other folate biosynthesis/metabolism genes. Since folate is essential for plant metabolism, it was surmised that the genes related to folate biosynthesis and metabolism are least likely to harbor genic polymorphism (Upadhyaya et al., 2017). Thus, the *FPGS* and *DHFS* knockout mutants led to embryo lethal phenotype in Arabidopsis (Mehrshahi et al., 2010). Even after double mutagenesis of tomato, it was found that genes involved in folate biosynthesis/metabolism were recalcitrant to mutagenesis (Gupta et al., 2017). The above reports highlighted that the folate levels in tomato are subtly modulated by cellular homeostasis (Upadhyaya et al., 2017).

Little is known about how the endogenous folate levels are determined by gene regulation, protein modifications, transport, and turnover. An analysis of QTLs in rice detected three QTLs upregulating folate levels in grains, but none was in folate biosynthesis genes (Wei et al., 2014). Likewise, two QTLs upregulated folate levels in maize, with a S-adenosyl-L- methionine-dependent methyltransferase and a transferase with the folic acid-binding domain as putative candidate genes (Guo et al., 2019). In tomato, mutations in the MYB117 transcription factor, namely the *tf-5* allele, upregulates the folate levels in fruits, indicating the role of transcription factors in determining the folate levels in plants (Tyagi et al., 2022).

The folate levels in plants may also be determined as part of cellular homeostasis by interaction with other metabolites and vitamins. In folate biosynthesis, pyridoxal 5′-phosphate (PLP, VitB6) is required as a cofactor for aminodeoxychorismate lyase (ADCL); thus, folate biosynthesis, in turn, may be determined by the levels of PLP in the plastids (Basset et al., 2004). Folate also controls redox homeostasis by assisting NADPH production, affecting cellular homeostasis (Gorelova et al., 2017b). Folate, a key constituent of one-carbon metabolism, provides C1 units to synthesize nucleic acid, pantothenate (vitamin B5), and amino acids in plants (Hanson and Roje, 2001).

Adding or removing one-carbon units is an important part of regulating plant metabolites levels and, in turn, affects the primary metabolism of plants. This is manifested by the linkage of folate with photorespiration, nitrogen metabolism (Jiang et al., 2013), biosynthesis of chlorophyll, betaine alkaloids, and lignin (Hanson and Roje, 2001), and the epigenetic regulation of flower transition in plants (Wang et al., 2017).

To comprehend how folate level is regulated in tomato and influences primary metabolism, we compared two contrasting tomato cultivars, Periyakulam-1 (PKM-1) and Arka Vikas (AV), bearing high folate and low folate levels in fruits, respectively. We report that high folate levels alter cellular homeostasis by elevating most amino acids. The high folate level in PKM-1 was also associated with the high carotenoid levels in PKM-1 fruit. We also show that the reduced activity of GGH in fruits is linked to increased folate levels in PKM-1 by higher folate polyglutamylation.

## Material and methods

### Plant materials and growth conditions

Tomato (*Solanum lycopersicum*) cultivar AV (Arka Vikas) and PKM-1 (Periyakulam- 1) were obtained from IIHR, Bangalore, India (www.iihr.res.in). The AV and PKM-1 were grown in the greenhouse as described in Bodanapu et al. (2016). The first two flowers from the first two trusses were tagged. The fruits were harvested at the mature green (MG), breaker (BR), and red-ripe (RR) stage. The fruits were homogenized in liquid nitrogen using a homogenizer (IKA, A11 basic, Germany), and the powder was stored at -80°C till further use.

### Biochemical analysis

The protocols used for different analyses were as follows: Gupta et al. (2014) for °Brix; Gupta et al. (2015) for carotenoids; Kilambi et al. (2013) for ethylene emission; Tyagi et al. (2015) for folate; GGH enzyme assay (Tyagi et al., 2022) and Bodanapu et al. (2016) for phytohormones and metabolites. Only metabolites with a ≥1.5-fold change (log2 ≥± 0.584) and P-value ≤0.05 were mapped on the metabolic pathway. Total RNA was extracted from the pericarp tissue of AV and PKM-1 fruits at different ripening stages using the hot phenol method (Verwoerd et al., 1989), and qRT-PCR was done as described previously by Kilambi et al. (2013). The qRT-PCR primers for folate and carotenoids biosynthesis pathway genes were designed using Primer 3 software (Table S1-2).

### Glutamylation, pABA, and pteridines extraction and quantifications

For quantifying glutamylation, all steps were similar to the folate extraction (Tyagi et al., 2015), except the step of rat plasma treatment was omitted. The pABA was extracted following Navarrete et al. (2012) protocol. Briefly, 100 mg powdered tissue was homogenized in 1 mL methanol and centrifuged at 2600*g* for 15 min at 4°C, the supernatant was collected, and the residue was re-extracted. The supernatants were pooled and dried by centrifugal evaporation. The dried sample was suspended in 1 mL Milli-Q water, followed by 5 min sonication. To 400 µL of the sample, 50 µL 2N HCl was added, and the mixture was incubated at 80°C for two hrs. The sample was cooled, and 2N NaOH was added to neutralize the acid. Lastly, the sample solutions were ultra-filtrated at 12000*g* for 12 min before LC-MS analysis, and 7.5 µL of the sample was injected into the column.

The pABA was separated on a reverse-phase Luna C18 column (5 µm particle size, 150 mm x 4.60 mm I.D.) (Phenomenex, CA, USA) using a gradient elution program. The gradient comprised of a binary solvent system consisting of 0.1% (v/v) of formic acid in water (solvent A) and methanol (solvent B) at a flow rate of 1 mL min^-1^. The injection volume was 7.5 µL, the run time was 15 min, and the column temperature was set at 24°C. The starting condition of the gradient was 92% of solvent A and 8% of solvent B. Subsequently, solvent B was linearly increased to 50% in 7 min, then to 100% in 10 min. After that, the mobile phase was reverted to the initial condition in 1 min and held for 4 min for re-equilibration of the column before the next injection. For all experiments, Exactive™Plus Orbitrap mass spectrometer (Thermo Fisher Scientific, MA, USA) was operated in the alternating full scan and equipped with positive heated electrospray ionization (ESI). The source parameters were optimized as follows: ion spray voltage, 4 kV; capillary temperature, 380°C; and heater temperature, 400°C. Sheath, auxiliary, and sweep gas flows were 75, 20, and 2 arbitrary units (au), respectively. For the full scan experiment, the mass scan was in the range of m/z 50-350. Instrument control and data processing were carried out by Xcalibur 3.0 software (Thermo Fisher Scientific). The pABA was detected in the positive mode with m/z 138.05 at the retention time of 5.0 min.

Pteridines were extracted following Martín-Tornero et al. (2016) protocol with some modifications. About 400 mg powdered tissue was homogenized in 1 mL methanol/water pH 12.0 (1/1, v/v) mixture, sonicated for 15 min, and centrifuged at 5000*g* for 10 min. The supernatant was collected, and the sample was re-extracted. Both supernatants were combined and dried by centrifugal evaporation. The dried residue was dissolved in 400 µL Milli-Q water. Lastly, 100 µL of sample solutions were ultra-filtrated at 12000*g* for 12 min before LC-MS analysis. The 7.5 µL sample was injected into the column.

Pteridines were separated using a gradient elution program on a reverse-phase Hypersil Gold C18 column (1.9 µm particle size, 50 mm x 2.1 mm I.D.) (Thermo Fisher Scientific). The gradient consists of a binary solvent system consisting of 0.1% (v/v) of formic acid in water (solvent A) and acetonitrile (solvent B) at a flow rate of 300 µL min^-1^. The injection volume was 7.5 µL, the run time was 8 min, and the column temperature was set at 24°C. For all experiments, Exactive™Plus Orbitrap mass spectrometer (Thermo Fisher Scientific) was operated in the alternating full scan and equipped with positive heated electrospray ionization (ESI). The source parameters were optimized as follows: ion spray voltage, 4 kV; capillary temperature, 380°C; and heater temperature, 320°C. Sheath, auxiliary, and sweep gas flows were 60, 20, and 2 arbitrary units (au), respectively. For the full scan experiment, the mass scan was in the range of m/z 50-500. Pteridines were detected in the positive ion mode with m/z for Npt (254.08), Mpt (254.08), HMPt, (194.06), and P6C (208.04) at the retention time of 2.63, 3.33, 5.57, and 5.68 respectively. Instrument control and data processing were carried out by Xcalibur 3.0 software (Thermo Fisher Scientific).

### PCA and metabolic pathway in AV and PKM-1

Principal Component Analysis (PCA) of primary metabolites was performed using Metaboanalyst 4.0 (http://www.metaboanalyst.ca/) after the One-way ANOVA plot (P ≤ 0.05) identified 66 of the 69 metabolites having significant contributions. The metabolites were mapped on the general metabolic pathways described in the KEGG (Kyoto Encyclopedia of Genes and Genomes, http://www.genome.jp/kegg) and LycoCyc (Sol Genomic networks, http://www.solcyc.solgenomics.net).

### Proteome profiling

Proteins (70 µg) were separated, identified, and quantified as described in Kilambi et al. (2016; 2021). *Solanum lycopersicum* iTAG4.1 proteome sequence (ftp://ftp.solgenomics.net/tomato_genome/annotation/ITAG4.1_release/ITAG4.1_proteins.fasta downloaded on August 28, 2020, 34430 sequences, and 11584472 residues) was used as the database against which the searches were done. The functional annotation and classification of the ITAG4.1 proteins was generated using the Mercator 4 tool (https://www.plabipd.de/portal/mercator4) on October 29, 2020 (http://www.plabipd.de/projects/Mercator4_output/GFA-62a085d6ce7e994eb0d871c612e7f528.zip) with the job id GFA- 62a085d6ce7e994eb0d871c612e7f528. The mass spectrometry proteomics data are available via ProteomeXchange (Vizcaíno et al., 2014) with the dataset identifier: Project Name: PKM- 1 proteome; Project accession: PXD027940. The proteins with ≥ 1.5-fold change (Log2 ± 0.58) and P-value ≤ 0.05 were considered as differentially expressed. For proteins detected only in PKM-1 or AV, the missing values were replaced with 1/10 of the lowest value in the dataset.

### Genome Sequencing

Whole Genome Sequencing and data analysis of AV and PKM-1 were done as described in Gupta et al. (2022). Briefly, sequencing was performed on the HiSeqX sequencing system (Illumina) (GeneWiz Inc. NJ, USA) according to the manufacturer’s protocol. A total of ∼200 million reads (31.5 Gb) with Q30 > 29 Gb data was generated. The raw reads were filtered using fastp software (v0.19.5) using parameters -M 30 -3 -5. The 2X 150 bp reads were mapped on *S. lycopersicum* cv. Heinz version SL3.0 using BWA-MEM (0.7.17) (Li and Durbin, 2009). The data analysis and variant calling was performed as described in Gupta et al. (2020). The resulting vcf (variant calling format) files were annotated using the SIFT4G algorithm. The effect of base substitutions on protein function was determined by SIFT4G (SIFT score ≤0.05 is considered deleterious) using the SIFT4G-ITAG3.2 genome reference database generated by Gupta et al. (2020).

### Statistical analysis

A minimum of three independent biological replicates was used for all experiments. Statistical Analysis On Microsoft Excel (https://prime.psc.riken.jp/MetabolomicsSoftware/StatisticalAnalysisOn-MicrosoftExcel) was used to obtain significant differences between AV and PKM-1. A Student’s *t*-test was also performed to determine significant differences (* for P ≤0.05). Heat maps 3D-PCA and graph plots were generated using Morpheus (https://software.broadinstitute.org/morpheus/) and MetaboAnalyst 4.0 (https://www.metaboanalyst.ca/), and SigmaPlot 11.0, respectively.

## Results

### Folate levels in PKM-1 and AV fruits

A large-scale screening of tomato accessions and cultivars revealed wide variation in folate levels of red-ripe fruits (Upadhyaya et al., 2017). To understand the molecular basis governing the variations in folate levels, we selected two tomato cultivars, Periyakulam-1 (PKM-1) and Arka Vikas (AV), with widely different folate levels. Both the cultivars are popularly grown by farmers in Southern India under similar agroclimatic conditions. Of these, PKM-1 displayed high (49.0±0.45 μg/100 g FW), and AV displayed low (21.2±1.72 μg/100 g FW) folate levels in red-ripe fruits (Fig. 1A) grown in the open field. Considering that the seasonal variations (Iniesta et al., 2009; Upadhyaya et al., 2017) influence folate levels, we reexamined the folate levels in PKM-1 and AV fruits in the next season. Though the folate level in the PKM-1 was slightly lower than in the earlier season, it was significantly higher than AV (PKM-1 40.4±1.7 μg/100 g FW; 18.1±2.0 μg/100 g FW) (Fig. 1A). As PKM-1 fruits had ≥2-fold higher folate levels than AV, notwithstanding seasonal differences, we compared these two cultivars to decipher the underlying basis of variations in folate levels. To minimize the influence of environmental factors on plant and fruit growth, we grew both PKM-1 and AV in the greenhouse for all following experiments.

**Fig. 1.**
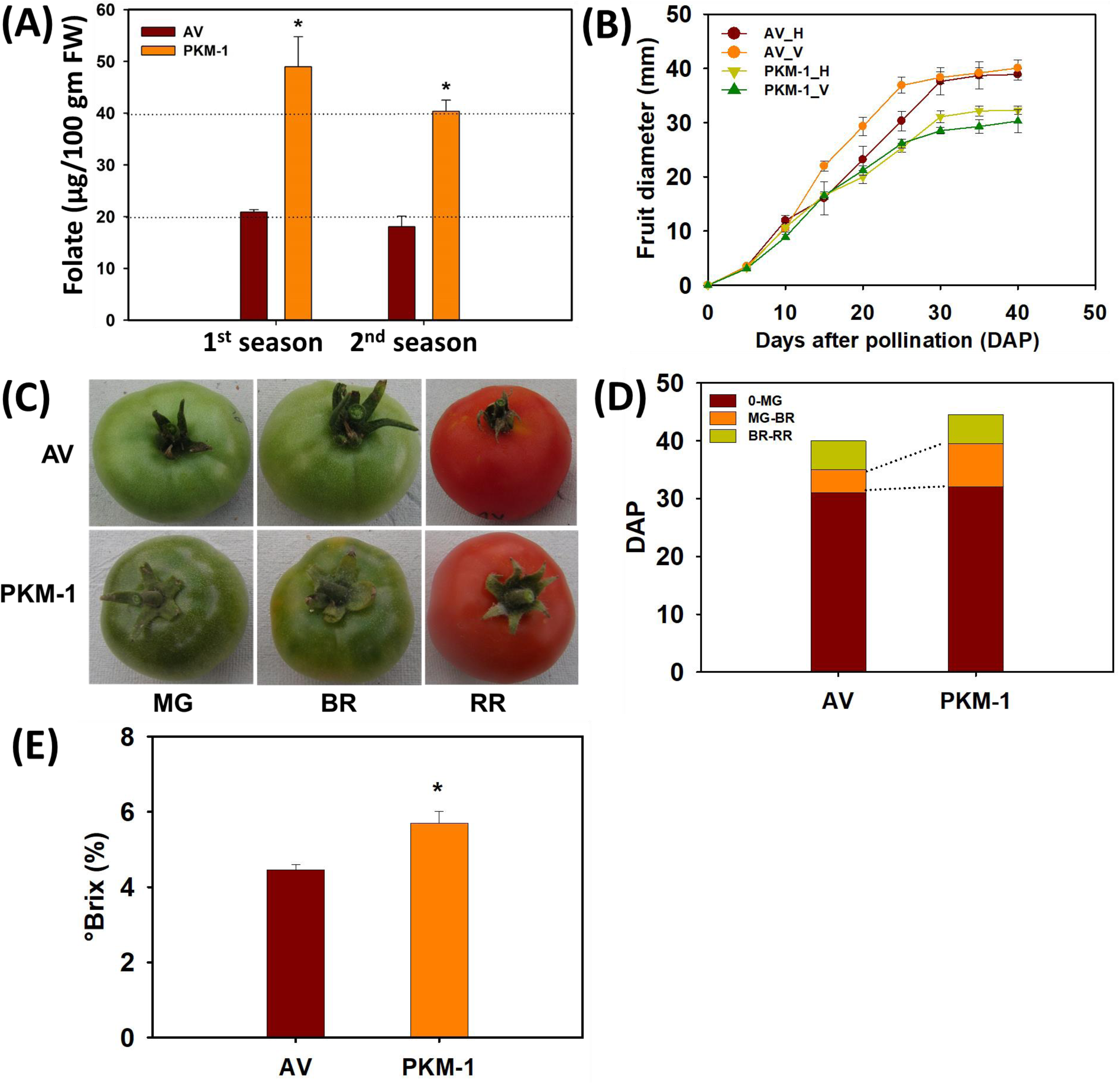
Folate and fruit ripening parameters of AV and PKM-1 fruits. (**A**) Folate content in 1^st^ and 2^nd^ season in PKM-1 at RR stage. (**B**) Time course of the increase in fruit diameter from anthesis until RR stage, (**C**) Fruit phenotype at mature green (MG), breaker (BR), and red-ripe (RR) stages of ripening, (**D**) Attainment of different ripening stages in tomato fruits. Note that PKM-1 is delayed in attaining in the BR stage. (**E**) °Brix in RR fruit. In (**A**), **(B)**, **(D)** and **(E)** data are means ± SE (n ≥ 3), * = P ≤ 0.05. An asterisk (*) shows a significant difference compared to AV. ***D***ays ***A***fter ***P***ollination.

### Fruit ripening in AV and PKM-1

The PKM-1 fruits were slightly smaller than AV; nonetheless, the on-vine growth of these fruits was nearly identical, reaching the near-final size 30 days after pollination (DAP) (Fig. 1B). Both AV and PKM-1 fruits reached the mature green (MG) stage at the same time (32 DAP) (Fig. 1C-D). However, post-MG stage PKM-1 fruits reached the breaker (BR) stage after seven days compared to 4 days for AV. Post-BR, both AV and PKM-1 fruits attained the red-ripe (RR) stage in 8-10 days (Fig. 1D). In tomato, ripening is also associated with increased sugar levels monitored as the °Brix value. The °Brix value in RR PKM-1 fruits was significantly higher than AV (Fig. 1E).

A major difference in PKM-1 and AV fruits was substantially lower ethylene emission from PKM-1 fruits (Fig. 2A). Both at BR and RR stage, the ethylene levels in PKM-1 were nearly 40% of AV. We then examined whether PKM-1 differed from AV regarding other phytohormones (Fig. 2B-H). Like ethylene, abscisic acid (ABA) level was nearly half in PKM- 1 at all ripening stages. PKM-1 also had a lower level of other phytohormones in a stage- specific fashion, jasmonic acid (JA) at MG, BR; salicylic acid (SA) at BR, RR; indole acetic acid (IAA) at BR. While indole-butyric acid (IBA) and methyl jasmonate (MeJA) did not vary significantly, RR fruits of PKM-1 had a higher level of zeatin at RR than AV.

**Fig. 2.**
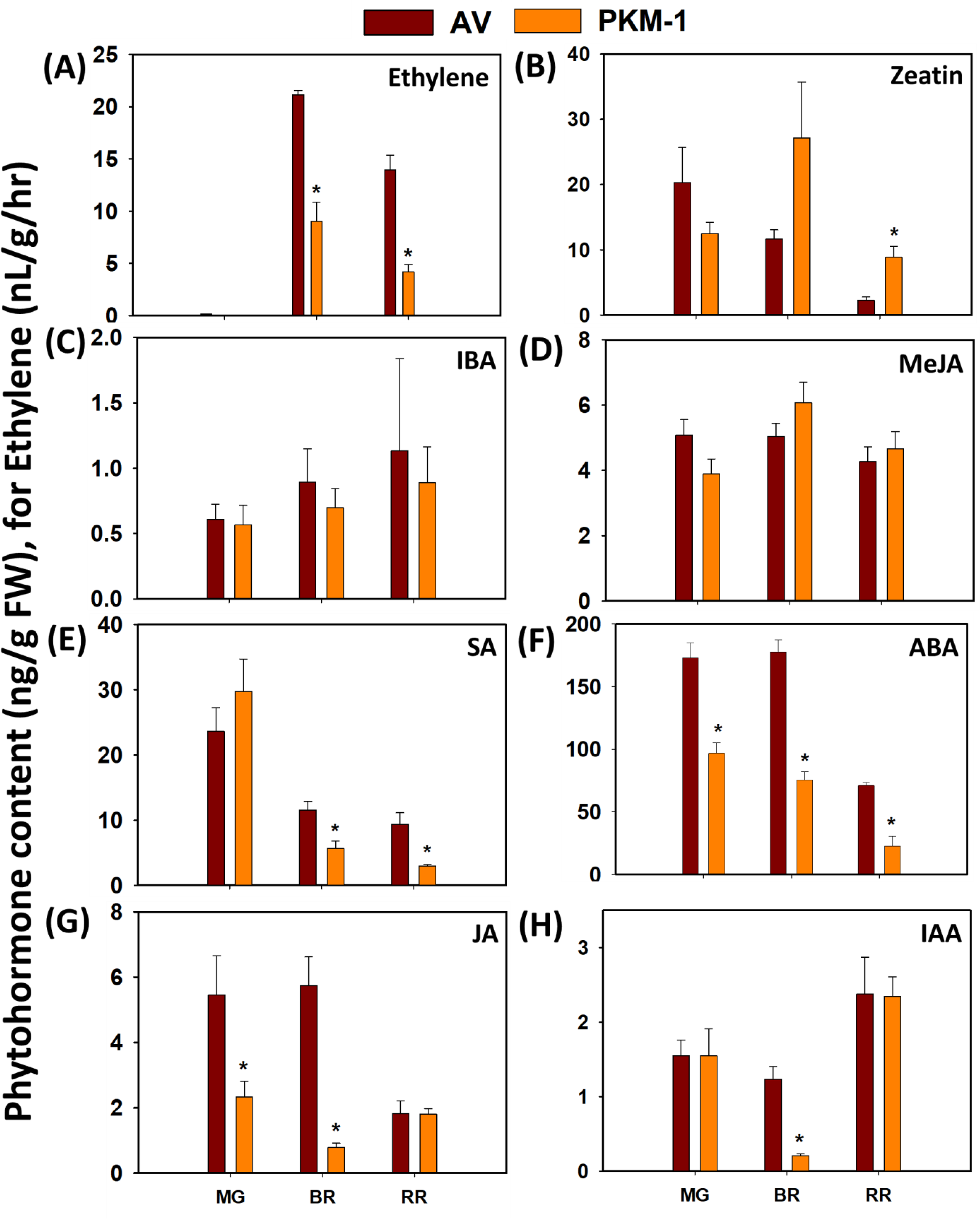
The level of phytohormones in AV and PKM-1 fruits during ripening. **(A)** Ethylene, (**B**) zeatin, (**C**) IBA, (**D**) MeJA, (**E**) SA, (**F**) ABA, (**G)** JA, and (**H)** IAA level in PKM-1. The levels of phytohormones were measured by LC-MS, except ethylene, which was measured by gas-chromatography and expressed as nL/g/hr. Data are means ± SE (n ≥ 3), * = P ≤ 0.05. An asterisk (*) shows a significant difference compared to AV. Abbreviations: IAA, indole acetic acid; IBA, indole butyric acid; MeJA, methyl jasmonate; SA, salicylic acid; ABA, abscisic acid, and JA, jasmonic acid.

### Folate levels during ripening of fruits

Though greenhouse-grown PKM-1 fruits had lower folate levels than the field-grown fruits (Fig. 1A), the folate level in PKM-1 was higher than AV at all ripening stages (Fig. 3A). During ripening, the folate level steadily declined, with the decline being higher in AV than in PKM-1, a pattern also seen in field-grown fruits. Among the four detected folate vitamers, viz. THF, 5-CH_3_-THF, 5,10-CH^+^THF, and 5-CHO-THF, THF were seen only at the MG at trace levels (Fig. S1 A-D). 5-CH_3_-THF was the major vitamer constituting 60-80% of folate, and 5,10-CH^+^THF and 5-CHO-THF contributed to the rest. The ripening-associated decline in total folate was solely contributed by 5-CH_3_-THF. The other two vitamers showed a marginal increase, particularly from MG to BR stage, barring 5-CHO-THF in AV. The 5- CHO-THF content was significantly high in PKM-1 than in AV at all stages of ripening.

**Fig. 3.**
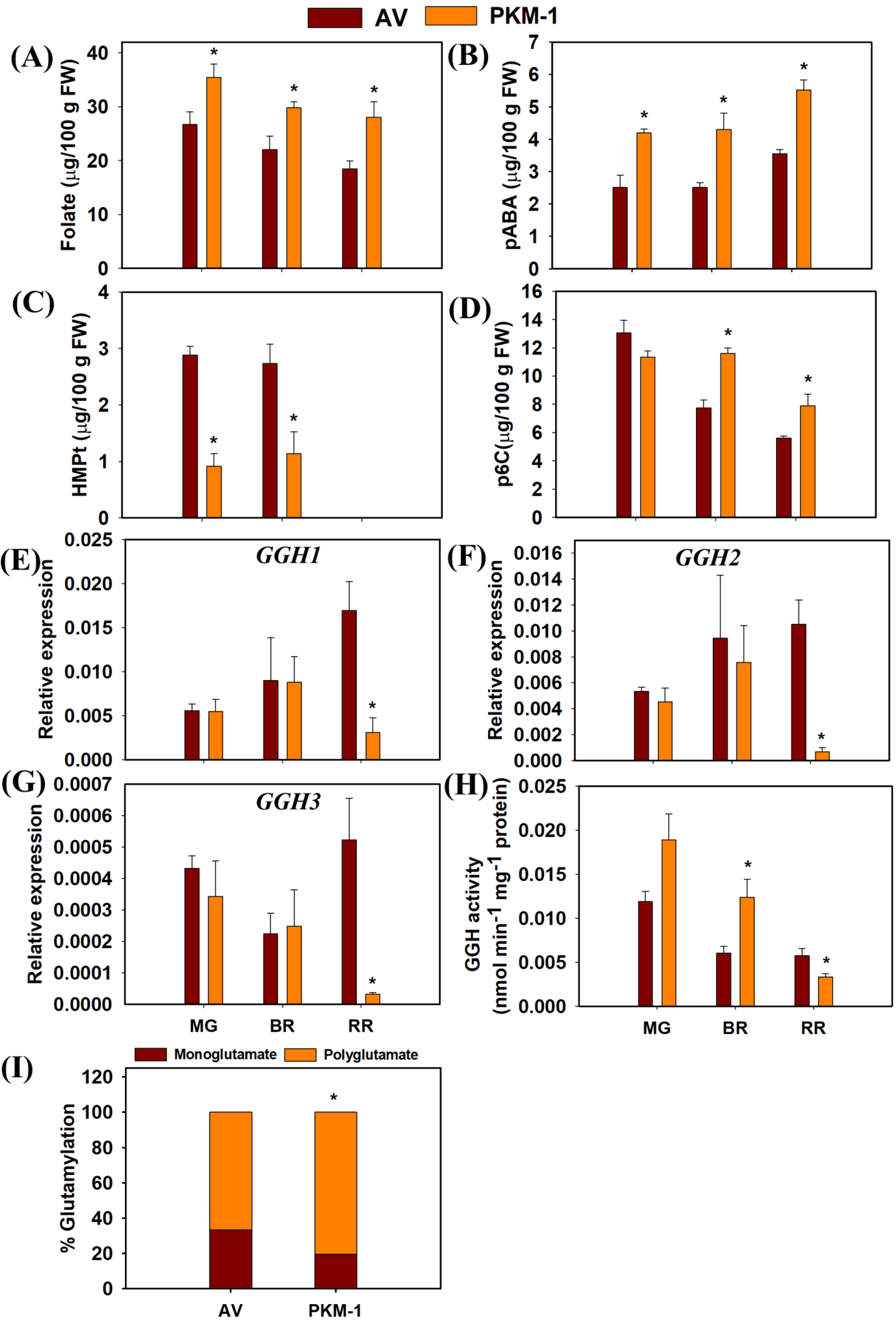
Folate, *p*ABA, folate salvage pathway intermediates, γ-glutamyl hydrolases (GGHs) transcript, activity, and level of glutamylation in PKM-1 fruit. (**A)** Folate, (**B**) p*-* aminobenzoic acid (*p*ABA), (**C**) 6-hydroxymethylpteridine (HMPt) and (**D**) pterin-6- carboxylate (p6C) levels in AV and PKM-1 at MG, BR, and RR stage. The level of (**E**) *GGH1*, (**F**) *GGH2*, and (**G**) *GGH3* transcripts in AV and PKM-1 fruits. (**H**) GGH enzyme activity in AV and PKM-1 fruits. (**I**) Percent polyglutamylation in AV and PKM-1 red ripe fruits. The reduced GGH activity is associated with higher polyglutamylation of folate. Data are means ± SE (n ≥ 3), * = P ≤ 0.05. An asterisk (*) shows a significant difference compared to AV.

We next examined whether the higher folate levels in PKM-1 are limited to fruits or seen in leaves. The PKM-1 leaves showed higher folate levels than AV (Fig. S2). The folate level in leaves was nearly 8-fold higher than in the RR fruits. In leaves too only four viz. THF, 5-CH_3_-THF, 5,10-CH+THF, and 5-CHO-THF folate vitamers were detected. In contrast to fruits, leaves showed a substantially higher level of THF.

### pABA levels are higher in PKM-1 fruits

We next examined whether higher folate levels in PKM-1 fruits arose due to increased precursors of folate biosynthesis, namely pABA and pterin. Parallel to higher folate levels, pABA was higher in PKM-1 fruits at all ripening stages. In AV and PKM-1, pABA levels increased from BR to RR stage (Fig. 3B). In contrast to pABA, we could not detect the pterins viz. 7-8-dihydroneopterin triphosphate and dihydroneopterin, the first and second intermediates of folate biosynthesis emanating from GTP. Presumably, their levels were lower than the detection limit.

Contrarily, we detected 6-hydroxymethylpterin (HMPt), an intermediate in the folate biosynthesis pathway and part of the folate salvage pathway. The other detected pteridine was 6-carboxypterin (p6C), a catabolite of pteridines and folates. Compared to AV, the level of HMPt was about 1/3rd in PKM-1 fruits at MG and BR (Fig. 3C). Remarkably, in both cultivars at the RR stage, the level of HMPt was below the detection limit. Contrasting to HMPt, the levels of p6C were nearly similar in PKM-1 and AV MG fruits. (Fig. 3D). In AV, the levels of p6C declined from MG to RR, while in PKM-1, the decline was from BR to RR stage.

### RR fruits of PKM-1 show lower GGH enzyme activity and higher polyglutamylation

Contrasting to higher folate levels, the transcript levels of most genes encoding for folate biosynthesis in PKM-1 were nearly similar to the AV (Fig. S3). The exceptions were *DHNA* at the BR stage, *DPP*, *DHFR,* and *HPPK*-*DHPS* at the RR stage, with lower transcript levels in PKM-1. The transcript levels of *DHFR* were high in PKM-1 at the MG stage (Fig. S3). Interestingly, the transcript levels of deglutamylation genes- *GGH1, GGH2,* and *GGH3* in PKM-1 were significantly lower than AV at the RR stage (Fig. 3E-G). The *in vitro* assay of the GGH enzyme showed that both in PKM-1 and AV, the activity of GGH was highest at the MG and declined with ripening progression (Fig. 3H). While GGH activity did not significantly differ at MG, at BR, GGH activity was significantly high in PKM-1, whereas at RR, the GGH activity was significantly low in PKM-1 fruits.

It is believed that polyglutamylated folate vitamers are less susceptible to catabolism than monoglutamylated forms. Also, folate-dependent enzymes prefer polyglutamylated folates over monoglutamate forms (Shane, 1989). The glutamylation status of folate vitamers was monitored with or without rat plasma conjugase at the RR stage to quantify the relative monoglutamate and polyglutamate levels. PKM-1 fruit has significantly higher polyglutamylation than the AV (Fig. 3I).

### RR fruits of PKM-1 show higher carotenoids levels

Since PKM-1 has higher folate levels, we checked whether the carotenoid levels were also affected in its fruits. Interestingly, PKM-1 RR fruits also had higher carotenoid levels than AV. Lycopene and β-carotene were 1.4 and 1.7-fold higher in PKM-1 fruits (Fig. 4). During ripening, the carotenoid profiling revealed a stage-specific upregulation in PKM-1, neoxanthin at MG and BR, violaxanthin at BR, δ-carotene at RR, and phytoene at RR and lutein at BR and RR stages (Table S3). Interestingly, γ-carotene was detected solely in RR PKM-1 fruits. While lycopene levels were very low at the MG and BR stage, β-carotene levels increased during ripening.

**Fig. 4.**
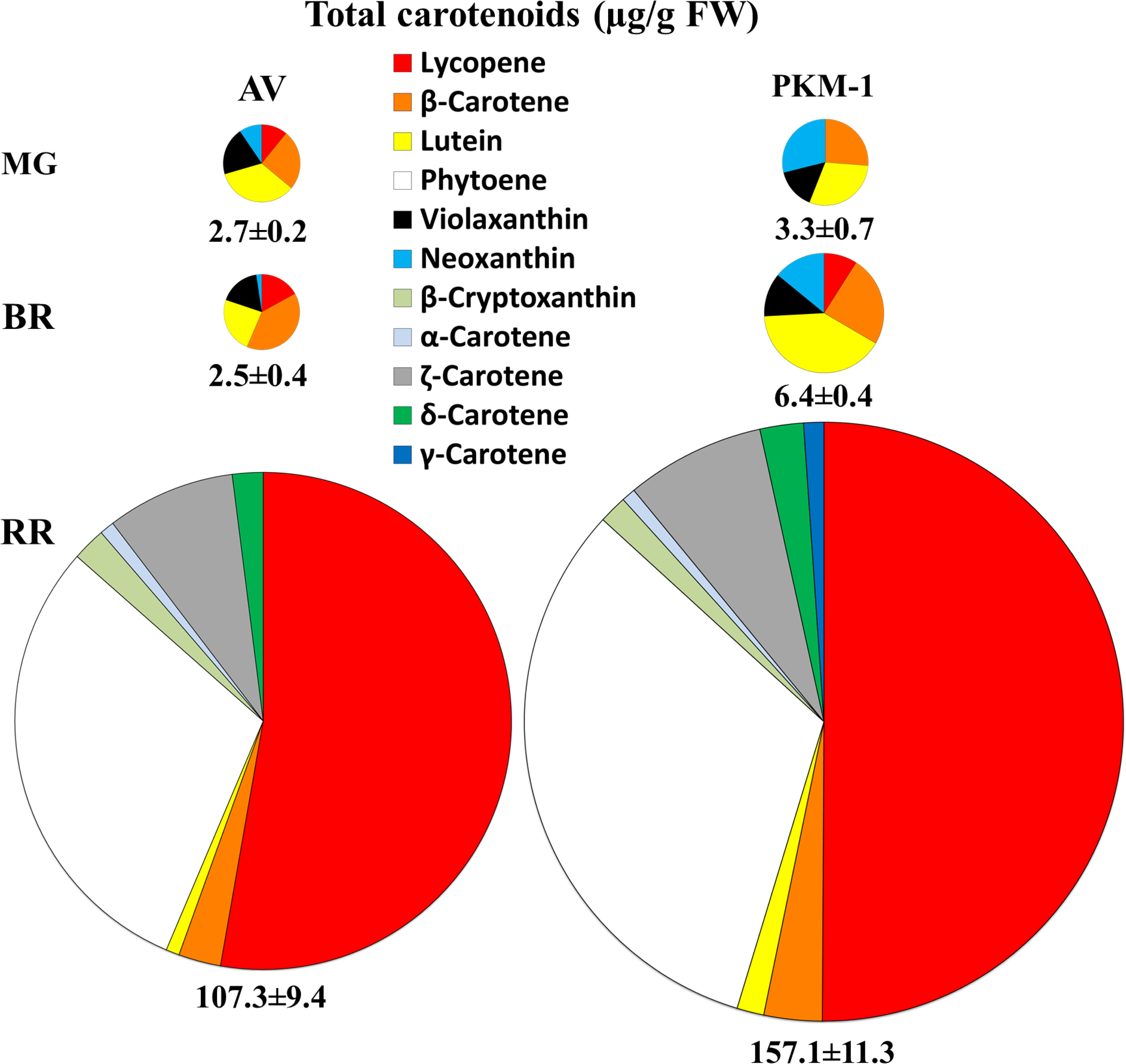
The proportional amounts of different carotenoids in AV and PKM-1 fruits at different ripening stages. The values adjacent to the individual pie indicate the total carotenoid content (μg/g FW). Note that the total area of the pie is proportional to the total carotenoid amount. The carotenoid data are means ± SE (n ≥ 4). See **Table S3** for individual carotenoid levels and significance.

We next analyzed the transcript levels of major genes involved in carotenoid biosynthesis (Fig. S4). The transcript levels of most genes were nearly similar between AV and PKM-1 at the MG and BR stages. Surprisingly, most genes contributing to lycopene formation

at the RR stage showed lower transcript levels in PKM-1 than AV, including *DXS*, *GGPPS2*, *PSY1*, *PDS*, *ZISO, ZDS, ZEP, NCED,* and *CYP97A* genes. At the MG stage, *CYCB* and *LCYB1* transcripts were high, whereas *PSY1*, *ZISO*, *NCED,* and *CYPC11* transcripts were low in PKM-1. At the BR stage, transcript levels of *PDS* and *LCYE* were significantly low in PKM-1.

### Metabolic profiles of PKM-1 fruits considerably differ from AV

Since AV and PKM-1 fruits markedly differed in folate level, a regulator of C1 metabolism, we examined whether metabolic profiles manifested similar differences. Out of 69 metabolites detected by GC-MS, most metabolite levels in PKM-1 substantially differed from AV. The metabolites included amino acids, amines, organic acids, fatty acids, sugars, and derivatives (Fig. 5A) (Dataset S1). The principal component analysis (PCA) displayed that at all stages of ripening, AV and PKM-1 metabolites were clustered separately (Fig. 5B). To delineate how changes in metabolite levels may alter cellular homeostasis, the significantly different metabolites of PKM-1 at different stages of ripening were mapped onto a general biochemical pathway (Fig. 5C).

**Fig. 5.**
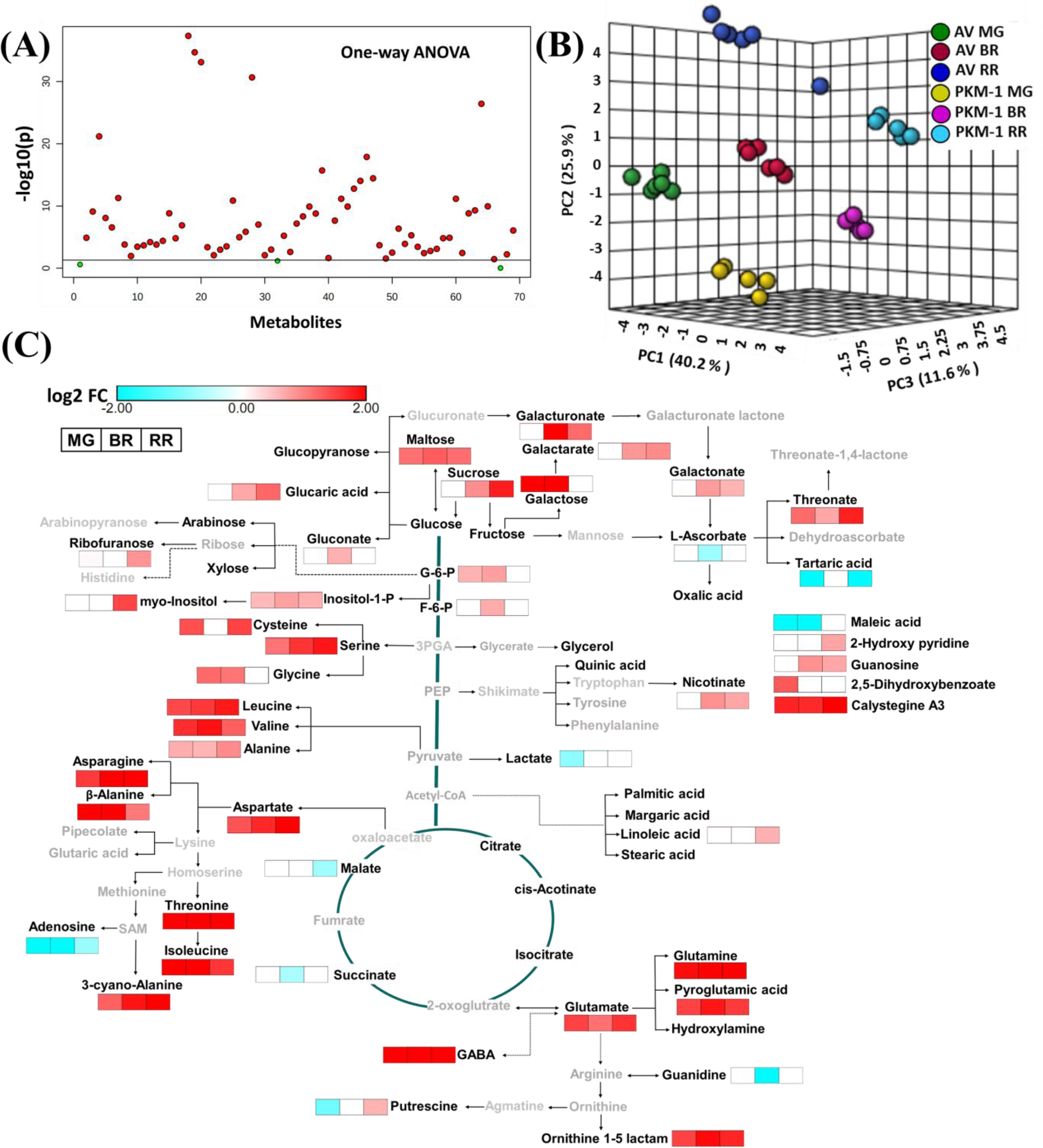
The levels of primary metabolites in AV and PKM-1 fruits at different ripening stages. (**A**) Important features (solid red circle) selected by a one-way ANOVA plot (P≤0.05). One-way ANOVA showed that 66 of the 69 metabolites present at MG, BR, and RR differed in AV and PKM-1. (**B**) Principal component analysis (PCA) of AV and PKM-1 at MG, BR, and RR stage. The PCA was plotted using MetaboAnalyst 4.0. (**C**) The metabolic shifts in PKM-1 fruits during ripening compared to AV. The relative changes in the metabolite levels at different ripening stages in PKM-1 fruits were determined by calculating the PKM-1/AV ratio at respective ripening phases. Only significantly changed metabolites are depicted on the metabolic pathway (log2 fold ≥ ± 0.584 and P-value ≤ 0.05). Bold black letters indicate the identified metabolites. Grey letters indicate metabolites below the detection limit. The white box indicates no significant change in metabolite level. Data are means ± SE (n ≥ 5), P ≤ 0.05. See **Dataset S1** for detailed metabolites data.

One distinct feature of the PKM-1 metabolome was elevated levels of most amino acids. Out of 20 L-amino acids, constituents of proteins, the levels of 13 amino acids were higher in PKM-1. In contrast to amino acids, sugars and sugar-derived metabolites showed a variable increase in PKM-1. In the glycolysis pathway, the glucose-6 phosphate (MG, BR) and fructose-6 phosphate (BR) were higher in PKM-1. Compared to sugars, TCA cycle intermediates- malate and succinate were lower at RR and BR stages, respectively.

### Proteome analysis in PKM-1 fruits

Considering the broad-spectrum effect of PKM-1 on carotenoids, folate, and metabolome, we profiled the proteome of AV and PKM-1 fruits. The proteome profiling of the AV and the PKM-1 revealed around 3309 (MG), 2940 (BR), and 2485 (RR) proteins in AV and 2570 (MG), 2383 (BR), and 2785 (RR) proteins in the PKM-1 cultivar (Fig. S5). Label- free quantification of proteins identified 493, 539, and 685 differentially expressed proteins (log2 fold ± 0.58, P-value ≤0.05) in PKM-1 at MG, BR, and RR, respectively. At MG and RR, most proteins were upregulated (270↑, 223↓, MG; 369↑, 316↓, RR), while in BR, most were downregulated (261↑, 278↓, BR) (Fig. 6A) (Dataset S2). The complement of up/down- regulated proteins varied in a stage-specific fashion, and only a few proteins overlapped at two or more stages.

**Fig. 6.**
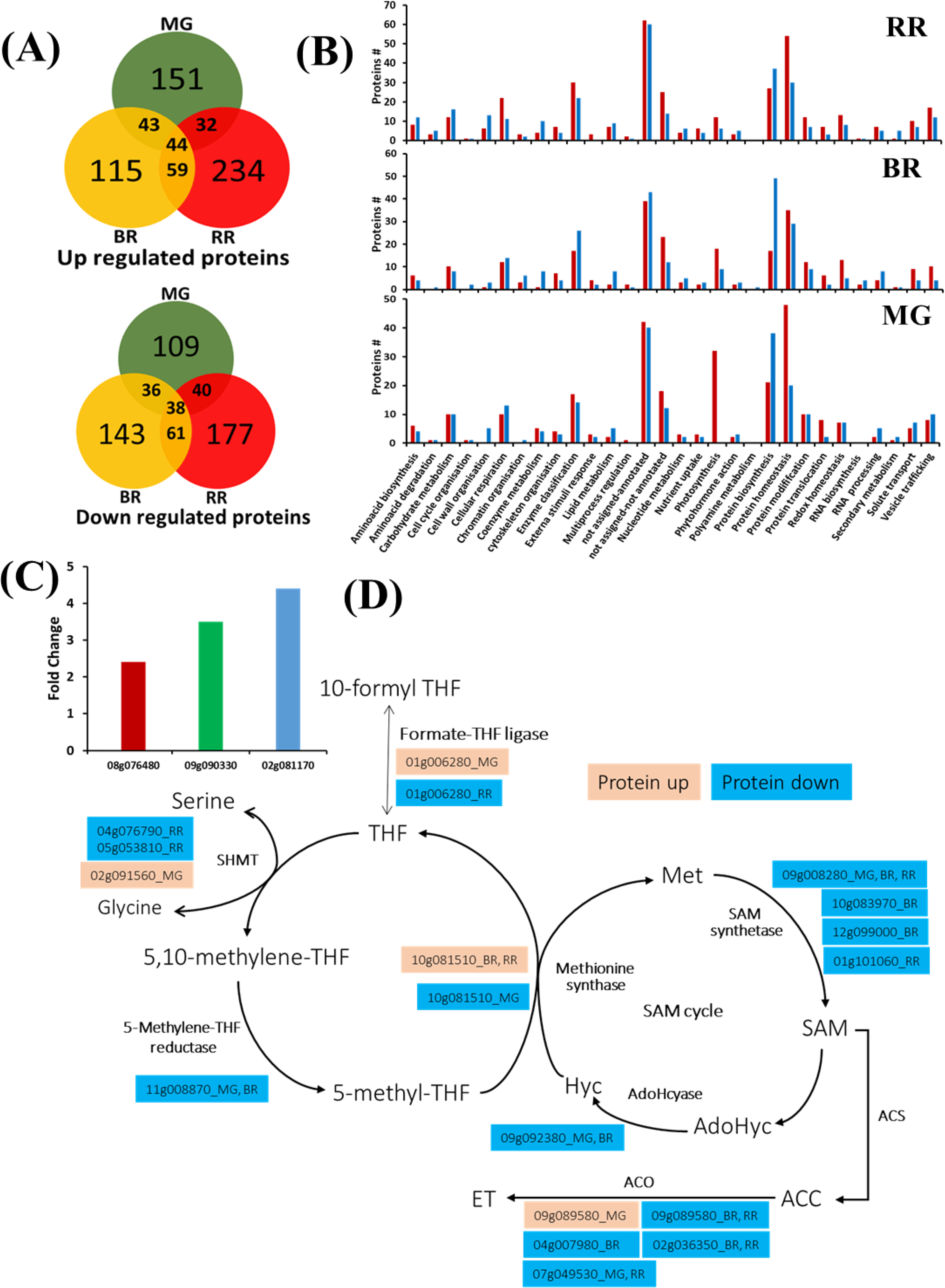
Proteome profiling, GO classification, carotenoid sequestration, and C1 metabolism and methionine pathway in PKM-1 and AV fruits at different ripening stages. **(A)** Differentially expressed upregulated and downregulated proteins in PKM-1 compared to AV (log2 fold change ≥ ± 0.58 and P ≤ 0.05). Note that only a few proteins are common between different ripening stages. **(B)** Functional classification of differentially expressed proteins in PKM-1 and AV. The differentially expressed proteins in the PKM-1 fruits compared to AV were functionally classified using Go-MapMan. **(C)** The fold change in the levels of carotenoids sequestering proteins such as harpin binding protein 1 (Solyc09g090330), CHRC (Solyc02g081170), and PAP3 (Solyc08g076480) in PKM-1. **(D)** C1 metabolism- related proteins in PKM-1 during ripening. Data are means ± SE (n=3), P ≤ 0.05. See **Dataset S2**. **MG**, Mature green; **BR**, Breaker; **RR**, Red-ripe.

The GO classification and Mapman analysis of differentially expressed proteins are in conformity with the wide-ranging influence of PKM-1 on cellular homeostasis (Fig. 6B, Fig. S5-6, Dataset S3). The largest classes of differentially expressed proteins belonged to protein homeostasis and biosynthesis. The higher °Brix in PKM-1 RR fruits correlated with upregulation of sucrose phosphate synthase (solyc07g007790). In PKM-1 fruits, the levels of carotenoids sequestering proteins such as harpin binding protein 1 (Solyc09g090330), CHRC, and PAP3 (Solyc02g081170, Solyc08g076480) were upregulated (Fig. 6C). In contrast, no specific accumulation of proteins leading to carotenoids biosynthesis or metabolism was discernible, except for the upregulation of MEP-pathway enzyme 4-hydroxy-3-methylbut-2- enyl diphosphate reductase (HDR) at RR.

Several proteins contributing to the primary precursors for amino acid biosynthesis were upregulated in PKM-1. The higher activity of aconitase (Solyc07g052350, Solyc12g005860 BR, RR) may direct TCA cycle flux towards glutamate, GABA (γ- aminobutyric acid), glutamine, and pyroglutamic acid. Likewise, flux through glycolysis is also directed towards higher amino acid levels. The high levels of phosphoglycerate kinase (Solyc07g066600 BR; Solyc07g066610 MG, BR, RR) may boost phosphoglyceric acid levels, the precursor for serine, glycine, and cysteine. The high levels of phoshopyruvate hydratase (Solyc09g009020, BR; Solyc10g085550 MG, BR, RR; Solyc03g114500 MG) may boost pyruvate levels leading to high levels of leucine, valine, and alanine. The level of 3- isopropylmalate dehydratase (Solyc06g060790), a protein in the leucine biosynthesis pathway, was upregulated in MG Conversely, γ-aminobutyrate transaminase I protein (Solyc07g043310) was downregulated in PKM-1 MG fruit.

The C1 metabolism was altered with ripening stage-specific expression of different components of methionine metabolism and folate transformations (Fig. 6D). Consistent with reduced ethylene emission from PKM-1 fruits, the levels of key ripening associated ethylene biosynthesis enzyme, 1-aminocyclopropane carboxylic acid synthase (ACO) 1 (Solyc07g049530), ACO6 (Solyc02g036350), and also of ripening associated protein E8 (Solyc09g089580) were downregulated (Fig. 6D). The reduced levels of S-adenosylmethionine synthase isoforms may have added to reduced ethylene emission.

### Genome Sequencing

To decipher the differences between AV and PKM-1, we sequenced and compared the whole genome of both cultivars. On the whole genome scale, PKM-1 had 350,316 SNPs that were not present in Arka Vikas, indicating that the PKM-1 genome differed by 0.0423% (Table S4). Notably, no polymorphic differences were found in genes encoding the folate and carotenoids biosynthesis pathways. The marking of deleterious mutations present in PKM-1 on tomato metabolic pathways revealed that only a few pathways were affected in PKM-1 (Fig. S7). Since most SNPs were localized in the noncoding regions of the PKM-1 genome, it is likely that the polymorphism in promoters of regulatory genes contributes to observed differences between Arka Vikas and PKM-1 (Meyer and Purugganan, 2013).

## Discussion

Tomato cultivars considerably differ in their °Brix, carotenoids, and folate levels (Mohan et al., 2016; Upadhyaya et al., 2017). Compared to AV, the PKM-1 had higher °Brix, carotenoids, and folate levels in RR fruits. Considering that little polymorphism exists in genes regulating carotenoids and folate levels, it was suggested that variation in their levels manifests interaction between transcriptome, proteome, and metabolome at cellular homeostasis level (Mohan et al., 2016; Upadhyaya et al., 2017). The comparative analysis of AV and PKM-1 is in conformity with the above notion for the regulation of °Brix, carotenoids, and folate levels.

### Several proteins modulating amino acid levels are altered in PKM-1

The paramount indicator of the alteration in cellular homeostasis is the levels of metabolites and proteins linked with glycolysis and the TCA cycle (Martínez-Reyes and Chandel, 2020; Skirycz et al., 2022). The higher °Brix and sucrose in PKM-1 fruits seem to be linked to increased sucrose synthase protein. While there were only a few stage-specific changes in levels of glycolysis and TCA cycle intermediates, a major change in PKM-1 was the elevation of a large number of amino acids barring those derived from the shikimate branch. In conformity with higher amino acid levels, two key proteins of the glycolysis pathway were upregulated in PKM-1. Among these, phosphoglycerate kinase provides precursor for serine, glycine, and cysteine, and phoshopyruvate hydratase provides a precursor for leucine, valine, and alanine.

Higher levels of aconitase, a TCA cycle protein, may be related to the elevation of glutamate, GABA, glutamine, and pyroglutamic acid. Additionally, the increased levels of glutamine synthase and glutamate synthase proteins may contribute to the above pathway. The reduced level of GABA transaminase I in PKM-1 may contribute to higher GABA levels, as the silencing of the above gene increases GABA in tomato fruits (Koike et al., 2013). Considering that these amino acids are derived from common intermediates in glycolysis and the TCA cycle, it is logical to assume that their higher levels in PKM-1 may relate to the upregulation of these proteins.

### Reduced ACO levels likely contribute to lower ethylene emission

One of the modulators of cellular homeostasis is hormones. The variations in hormonal levels have a cascading influence on plant metabolome and development (Fàbregas and Fernie, 2021). The lower levels of ABA, SA, and MeJA indicate a major alteration in the hormonal profile of PKM-1 fruits. In tomato, ethylene is needed for fruit ripening, as the block in ethylene perception or biosynthesis negatively affects ripening (Sharma et al., 2021). The PKM-1 fruits emitted substantially lower ethylene than AV, which seems to be related to lower amounts of ACO1 and ACO6, two key proteins contributing to ripening-specific ethylene synthesis (Houben and Van de Poel, 2019). The reduced ABA, SA, and JA levels seem to be related to reduced ethylene emission in PKM-1. A similar lowering of ABA, SA, and JA is seen in the *acs2-2* mutant compromised in ethylene biosynthesis (Sharma et al., 2021).

It can be construed that the alteration of the hormonal profile in PKM-1 may be responsible for the observed metabolic differences between PKM-1 and AV. Nonetheless, the linkage between a metabolite and hormones and developmental response is complex (Fàbregas and Fernie, 2021). In tomato, the RNAi-suppression of invertase (LIN5) reduces °Brix levels, leads to smaller fruits, and lowers sucrose synthase activity and levels of ABA and JA (Zanor et al., 2009). Contrarily, in PKM-1, higher °Brix is associated with increased sucrose synthase protein but reduced ABA and JA levels. Despite lower ethylene emission, the transition period of PKM-1 fruits from the MG to RR stage was only moderately longer. Probably, the ethylene emission from PKM-1 fruit was still sufficient; therefore, ripening duration remained unaffected.

### Increased sequestration may be responsible for the elevation of carotenoids levels

The upregulation of carotenoids during tomato ripening is influenced by many causative factors with a complex web of interactions (Liu et al., 2015). Among these factors, ethylene is a major factor in modulating fruit-specific carotenogenesis. The transgenic suppression of ethylene biosynthesis or loss of ethylene perception in *Nr* mutant inhibits carotenoid accumulation (Theologis et al., 1993; Lanahan et al., 1994). Besides ethylene, ABA and JA also stimulate carotenoid accumulation in tomato (Liu et al., 2015). Considering that PKM-1 fruits, despite having reduced levels of ethylene, ABA, and JA, accumulate higher levels of carotenoids, the upregulation of carotenoids seems unrelated to the hormonal profile of fruits. Likewise, the tomato *Nps1* mutant has higher carotenoid levels but reduced emission of ethylene and ABA levels (Kilambi et al., 2021).

The upregulation of carotenoids level in tomato fruits has also been ascribed to increased expression of carotenoid biosynthesis pathway genes, particularly phytoene synthase 1 (Fraser et al., 2007). The lower or nearly similar transcript level of most carotenoid biosynthesis genes in PKM-1 indicates that the stimulation of carotenogenesis is not related to transcript levels. The possibility that reduced transformation of carotenoids to xanthophylls may boost lycopene levels is unlikely, as there was no reduction in xanthophylls in PKM-1.

The enhanced level of carotenoids in PKM-1 fruits seems to be more related to efficient sequestration by carotenoids binding proteins (Vishnevetsky et al., 1999). Emerging evidences support that fibrillin family proteins enable a higher accumulation of carotenoids in chromoplasts. In tomato, high pigment mutants increase CHRC protein, which likely boosts carotenoid levels by sequestration (Kilambi et al., 2013). The overexpression of pepper fibrillin increases carotenoid levels in tomato fruits (Simkin et al., 2007). Contrarily, VIGS of fibrillins reduce lycopene accumulation in tomato fruits (Pourcq, 2014). In pepper, the harpin binding protein assists the formation of lycopene crystals (Deruère et al., 1994). The increased level of lycopene in PKM-1 may be related to increased levels of carotenoid sequestration proteins, viz., harpin binding protein 1, and CHRC and PAP3. Ostensibly, an increase in carotenoid sequestration proteins seems to be the causative factor for higher lycopene in PKM-1 fruits rather than modulation of carotenoid biosynthesis or degradation,

### Elevation of folate mildly affects C1 metabolism in PKM-1 fruits

The interrelationship between folate levels and tomato ripening seems to be more complex. In PKM-1 and AV, post-MG THF was not detectable, perhaps due to lower folate biosynthesis or faster conversion of THF to other vitamers. In most tomato cultivars, folate levels decline during ripening, and only a few cultivars show increase during the ripening (Upadhyaya et al., 2017). In PKM-1 and AV, total folate levels declined during ripening, albeit in PKM-1, the decline was slower. Among three-folate vitamers detected in ripening fruits, the decline was confined to 5-methyl-THF, the major vitamers in leaf and fruit. The reduction in 5-methyl-THF may be related to the increased levels of methionine synthase in PKM-1. Additionally, variable expressions of MTHFR, SHMT, and formate-tetrahydrofolate ligase were observed at different ripening stages. Taken together, the elevated folate levels in PKM- 1 only mildly influenced the C1 metabolism.

Notwithstanding the above, the emerging evidence has brought forth a novel role for 5- CHO-THF in plants. So far, the CHO-THF is considered an intermediate in the C1 metabolism, which does not serve as a C1 donor. Rather it binds to SHMT and other folate-dependent enzymes and inhibits their enzymic activity, at least *in vitro* (Stover and Schirch, 1993). Recent evidence indicated that 5-CHO-THF binds to many proteins and can modulate C/N metabolism (Li et al., 2021). Considering that 5-CHO-THF levels are higher in PKM-1, it may influence C1 metabolism and the C/N metabolism, signified by elevated amino acid levels in PKM-1.

### Elevated folate levels in PKM-1 are not related to folate biosynthesis

The higher folate level in PKM-1 may reflect a higher synthesis rate or reduced degradation of folate vitamers. Between the two precursors of folate biosynthesis, the pterin was below the detection limit. In contrast, the PABA level was higher in ripening PKM-1 fruit. The transgenic manipulation of pterin and PABA biosynthesis in tomato fruits indicated that a combined upregulation of both precursors is needed for increased folate levels. The transgenic upregulation of pterin led to a 2-fold increase in folate, while upregulation of PABA did not influence the total folate level (de la Garza et al., 2007). Since the pterin level was below detection limits, whether a higher PABA level in PKM-1 fruit elevated the total folate level remains ambiguous.

In tomato, including PKM-1, the leaves have several-fold higher folate levels than fruits (Tyagi et al., 2015). It can be construed that folate biosynthesis is attenuated in fruits than leaves. Consistent with this, the transcript levels of key folate biosynthesis genes *ADCS* and *GCHI* decline during tomato ripening, and other genes show variable expression (Waller et al., 2010). In PKM-1, like in maize (Lian et al., 2015), transcript levels of folate biosynthesis genes also show variable expression at different ripening stages and thus had little correlation with folate biosynthesis. Ostensibly, high folate level in PKM-1 seems to be mediated by a process different from the regulation of folate biosynthesis pathway genes.

### Reduced GGH activity likely contributes to higher folate levels in PKM-1 fruits

In addition to biosynthesis, the endogenous folate level is also determined by degradation and the extent of polyglutamylation. In plants, the rate of folate breakdown is ca. 10% per day (Strålsjö et al., 2003). The non-enzymatic cleavage of the C9-N10 bond in folate yields dihydropterin-6-aldehyde and pABA-Glutamate. As folate is important for metabolism, plants have a folate salvage pathway that regenerates pterin and *p*ABA from these breakdown products. The reduced levels of 6-hydroxymethylpteridine (HMPt) in PKM-1 may be due to its faster conversion to folate precursor (Gorelova et al., 2017a). Alternately, the pterin aldehyde reductase (PTAR) in PKM-1 may be less efficient, as manifested by higher levels of pterin-6-carboxylate in ripening fruits. Together, the above two processes may determine the extent of recycling the pterin to the folate biosynthesis pathway.

The in vivo folate level can be altered by shortening or lengthening polyglutamyl tails. The relative activities of FPGS, which adds glutamate tail to folate, determine the extent of folate polyglutamylation. Conversely, the activities of GGHs that remove glutamate tail also influence the *in vivo* folate levels. In tomato, the overexpression of *GGH* resulted in extensive deglutamylation of folate and lowered the total folate by 40% (Akhtar et al. 2010). Considering that in PKM-1, the polyglutamate percentage in folate is significantly high, indicating reduced activity of GGH. In conformity with this, PKM-1 has lower *GGH1*, *GGH2,* and *GGH3* expressions and reduced GGH enzyme activity. Similarly, in tomato *tf-5* mutant, the increased folate levels in fruits strongly correlated with the reduced GGH activity (Tyagi et al., 2022). Taken together, the high folate levels in PKM-1 seem to be related to lowering the deglutamylation process as it has lower GGH activity in fruits.

While there were polymorphic differences between both cultivars’ genome sequences, only 830 genes in PKM-1 bore SNPs, possibly leading to loss of function (Table S4, Dataset S4). Since most of above genes belong to large gene families, their loss may be offset by other members of family. The genome sequence comparison highlighted that observed differences in carotenoids and folate levels in these cultivars are not due to polymorphic differences in genes regulating folate and carotenoids biosynthesis. Since majority of SNPs resides in noncoding region, it is reasonable to assume that the diversity of metabolic changes in PKM-1 compared to AV ensues from polymorphic differences in promoter regions. In potato, out of 497 SNPs influencing folate level, a lone SNP was located within the 5-formyltetrahydrofolate cycloligase gene, and only 17 SNPs were in proximity of promoter/enhancer regions of folate metabolism genes (Bali et al., 2018). The SNPs in promoters can affect gene expression by modifying *cis*-regulatory elements and binding of transcription factors, as indicated for *acs2-2* mutant in tomato (Sharma et al., 2021). Broadly, polymorphic differences in promoters may influence transcriptional regulations of genes leading to alteration of cellular homeostasis including higher folate and carotenoid levels in fruits.

To sum, our study shows that PKM-1 RR fruits have elevated levels of three nutraceuticals, carotenoids, GABA, and folate. The metabolome and proteome profiling also divulge the diversity in regulating these three nutraceuticals in PKM-1 fruits. The carotenoid sequestration proteins seem to be a key regulator for increased carotenoid levels. Conversely, lowering of degradation of GABA and reduced deglutamylation of folate seems to be key loci for higher GABA and folate, respectively, in tomato. In recent years, GABA has been promoted as a health-promoting nutraceutical as exemplified by approval for commercial cultivation of a gene-edited tomato enriched in GABA in Japan (Waltz, 2022). In conclusion, this study provides the potential gene targets for improving carotenoids, GABA, and folate in tomato.

## Acknowledgments

This work was supported by the Department of Biotechnology (DBT), India grants, BT/PR11671/PBD/16/828/2008, BT/PR/7002/PBD/16/1009/2012, and BT/COE/34/SP15209/2015 to RS and YS. We also acknowledge fellowships by CSIR to KT and UGC to AS and HVK.

## Author contributions

KT, YS, and RS designed the research and wrote the manuscript; KT did most experiments, AS did proteome data analysis, HVK did the proteome profiling. PG analyzed the AV and PKM-1 genome sequence.

## Data Availability

The data consisting of metabolite levels and proteome is included as supplemental material. The proteome data are uploaded to the PRIDE repository, which can be accessed using the following details: Project Name: PKM-1 proteome; Project accession: PXD027940.

## Supplementary information

Additional Supporting Information may be found online in the Supporting Information tab for this article.

**Table S1.** List of primers used for qRT-PCR analysis for folate biosynthesis pathway.

**Table S2.** List of primers used for qRT-PCR analysis for carotenoid biosynthesis pathway.

**Table S3.** Carotenoids profiles of AV and PKM-1 fruits at MG, BR, and RR stages.

**Table S4.** The relative proportions of different functional categories of mutations in genes and in noncoding regions, including promoters and introns.

**Dataset S1.** Metabolites profile of PKM-1 and its WT during fruit ripening.

**Dataset S2.** Proteome profile of PKM-1 and its WT during fruit ripening.

**Dataset S3.** Differentially expressed proteins of *tf-5* mentioned in text/Figures.

**Dataset S4.** Single nucleotide polymorphism (SNPs) identified through Whole genome sequencing in PKM-1.

---

**Fig. S1.**
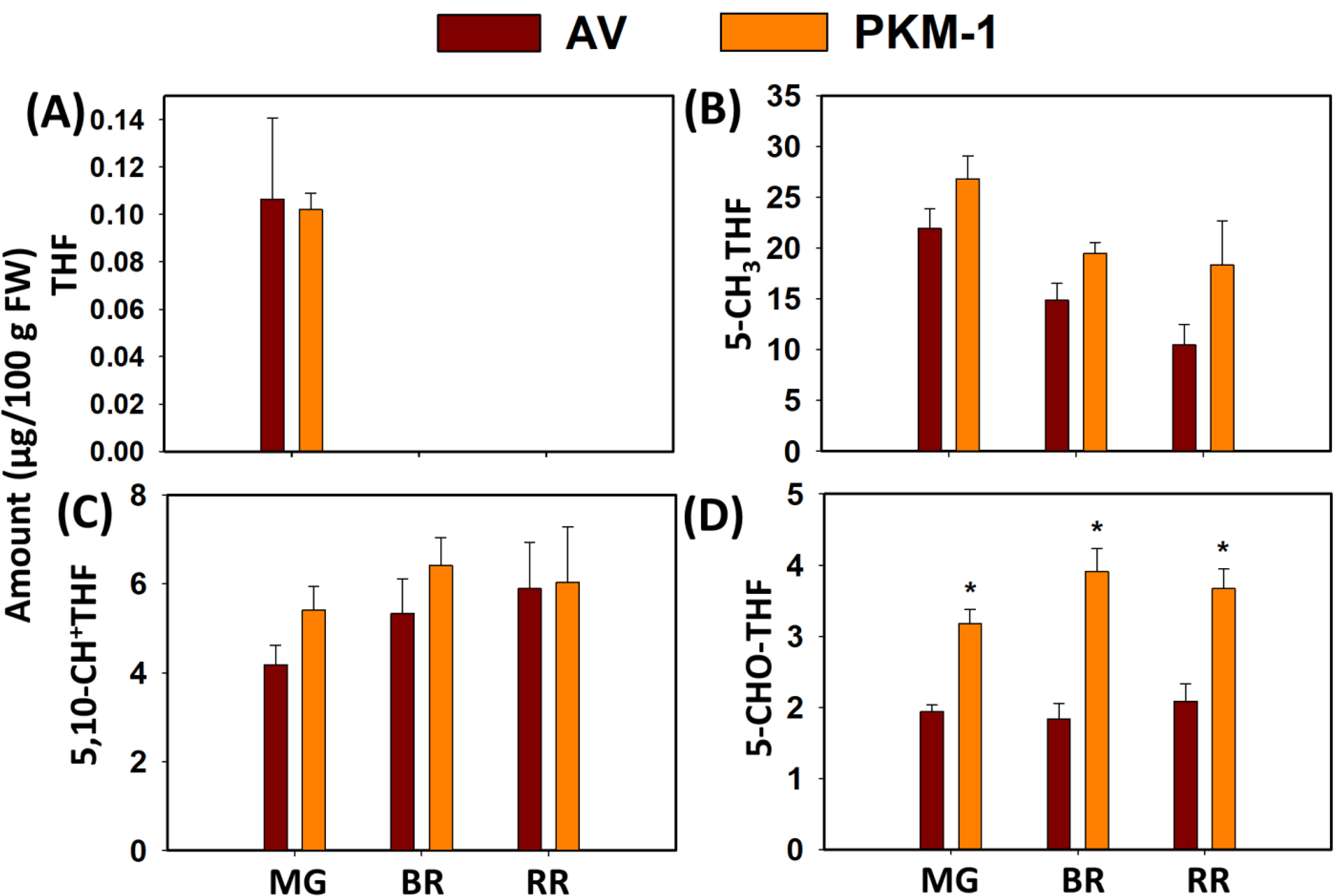
Folate profiling in AV and PKM-1 during ripening. (**A**) THF, (**B**) 5-CH_3_THF, (**C**) 5,10-CH^+^THF, and **(D)** 5-CHO-THF at different ripening stages. Data are means ± SE (n ≥ 3), * = P ≤ 0.05. An asterisk (*) shows a significant difference compared to AV. Abbreviations: THF, tetrahydrofolate; 5-CH_3_THF, 5-methyltetrahydrofolate; 5,10-CH^+^THF, 5,10- methenyltetrahydrofolate; 5-CHO-THF, 5-Formyltetrahydrofolate.

**Fig. S2.**
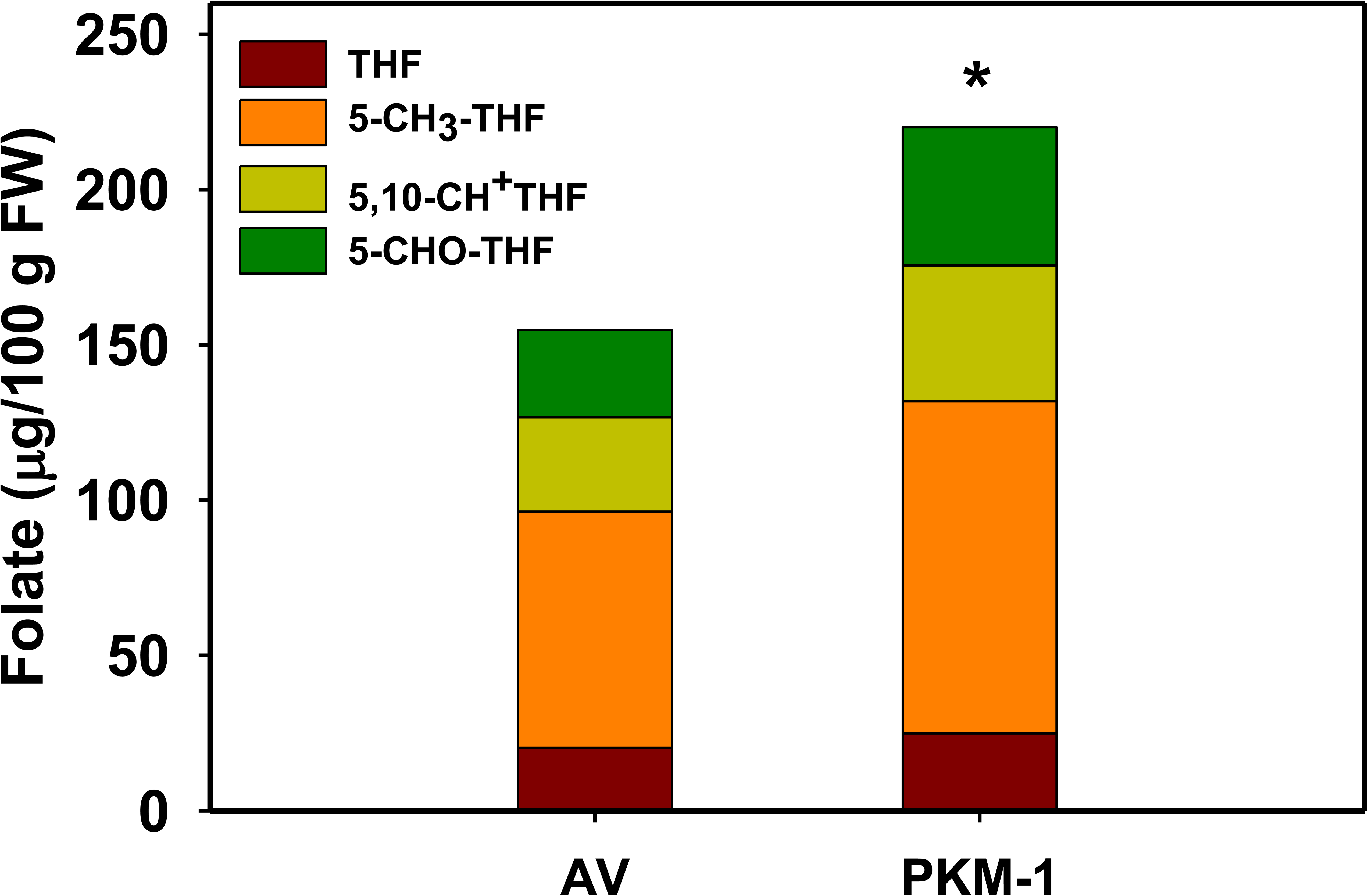
Total folate levels in AV and PKM-1 leaves. The leaves harvested from the 5^th^ node of 7-week-old AV and PKM-1 plants were used for folate estimation using LC-MS. Data are means ± SE (n = 5), * = P ≤ 0.05. An asterisk (*) shows a significant difference compared to AV Abbreviations: THF, tetrahydrofolate; 5-CH_3_THF, 5-methyltetrahydrofolate; 5,10- CH^+^THF, 5,10-methenyltetrahydrofolate; 5-CHO-THF, 5-Formyltetrahydrofolate.

**Fig. S3.**
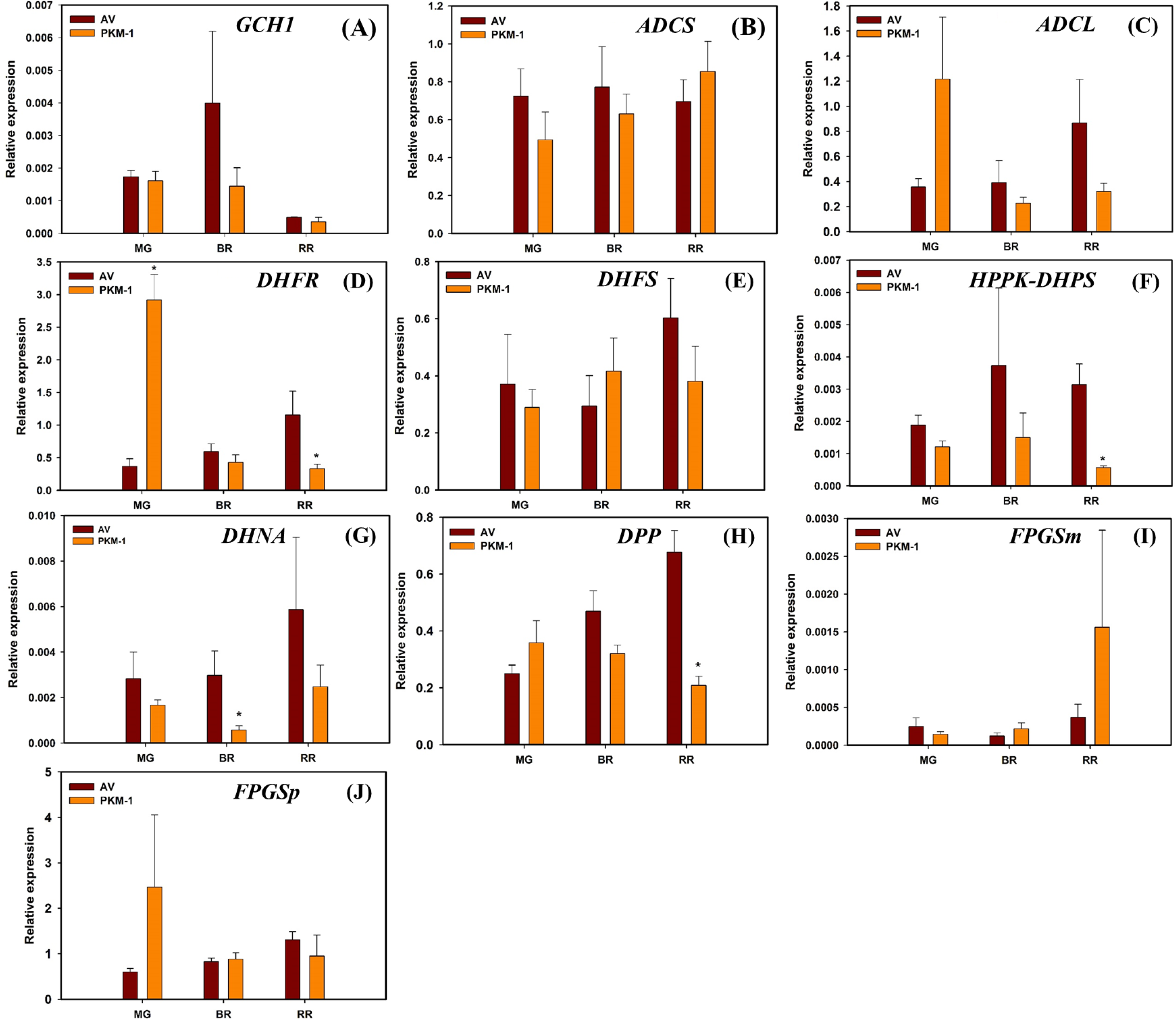
The expression of folate biosynthesis pathway genes in AV and PKM-1 fruits. Data are means ± SE (n = 3), * = P ≤ 0.05. An asterisk (*) shows a significant difference compared to AV Abbreviations: *GCH1*, GTP cyclohydrolase I; *DPP*, dihydroneopterin (DHN) triphosphate diphosphatase; *DHNA*, DHN aldolase; *ADCS*, aminodeoxychorismate (ADC) synthase; *ADCL*, aminodeoxychorismate lyase; *HPPK-DHPS*, 6-hydroxymethyl-7,8- dihydropterin (HMDHP) pyrophosphokinase (HPPK) and dihydropteroate (DHP) synthase; *DHFS*, dihydrofolate (DHF) synthase; *DHFR*, DHF reductase; *FPGS*, folylpolyglutamate synthase (p-plastic, m-mitochondrial).

**Fig. S4.**
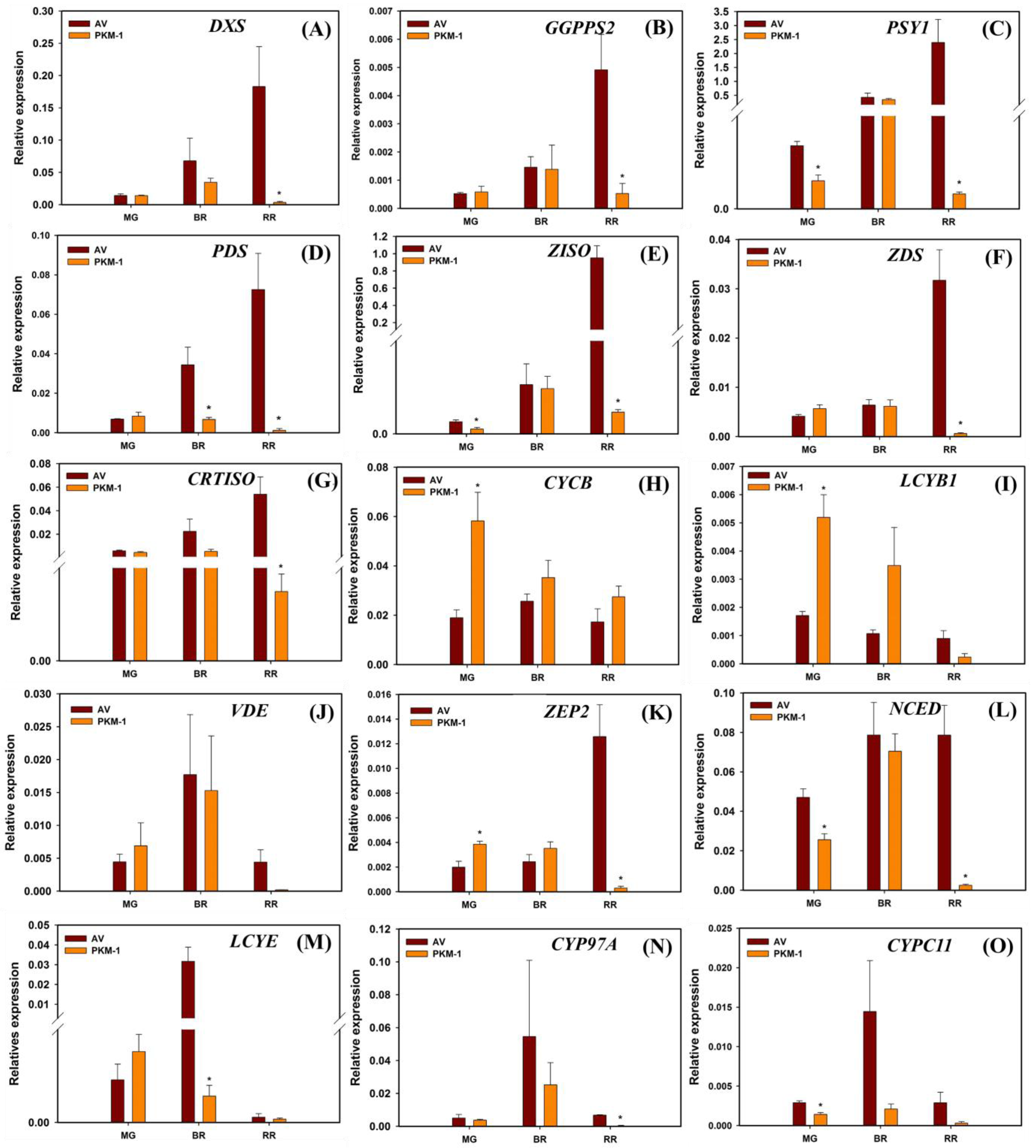
Carotenoid biosynthesis pathway gene expression in AV and PKM-1 fruit during ripening. Data are means ± SE (n = 3), * = P ≤ 0.05. Abbreviations: *DXS*, 1-deoxy-d-xylulose- 5-phosphate synthase; *GGPPS2*, geranylgeranyl diphosphate synthase; *PSY1*, phytoene synthase; *PDS*, phytoene desaturase; *ZISO*, ζ-carotene isomerase; *ZDS*, ζ-carotene desaturase; *CRTISO*, carotenoid isomerase; *CYCB*, lycopene β-cyclase (chromoplastic); *LCYB1*, lycopene β-cyclase; *VDE*, violaxanthin de-epoxidase, *ZEP*, zeaxanthin epoxidase; *NCED*, 9-cis- epoxycarotenoid dioxygenase; *LCYE*, lycopene ε-cyclase, *CYP97A*, cytochrome P450 carotenoid β-hydroxylase; *CYPC11*, cytochrome P450 carotenoid ε-hydroxylase.

**Fig. S5.**
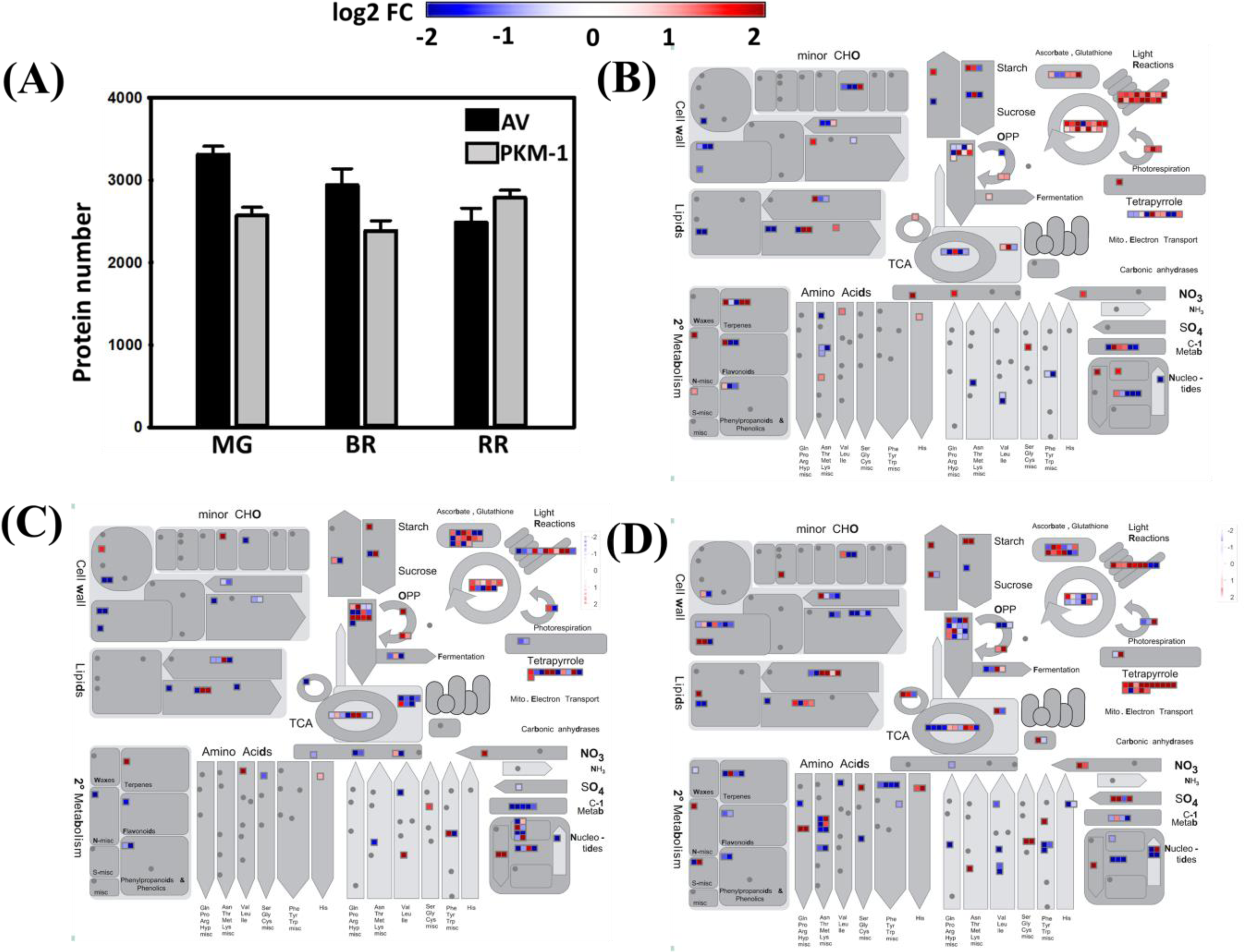
Total protein identified in AV and PKM-1 and Mapman representations of proteome changes in PKM-1 fruits. **(A)** Proteins were identified in AV and PKM-1 at MG BR and RR stage through Sorcerer software version 1. (**B-D)** Mapman overview of metabolic processes affected in PKM-1 at different stages of fruit ripening. **B-D** shows processes affected by differentially expressed proteins at MG **(B)**, BR **(C),** and RR **(D)** stages. Each square represents one protein. The red square indicates the upregulation of transcripts/proteins, and the blue square indicates the downregulation of transcripts/proteins. Only significantly different proteins (log2 fold ≥±0.58, P ≤0.05) are depicted in heat maps. Data are means ± SE (n = 3), P ≤ 0.05. (For details, see **Dataset 3). MG**, Mature green; **BR**, Breaker; **RR**, Red-ripe.

**Fig. S6.**
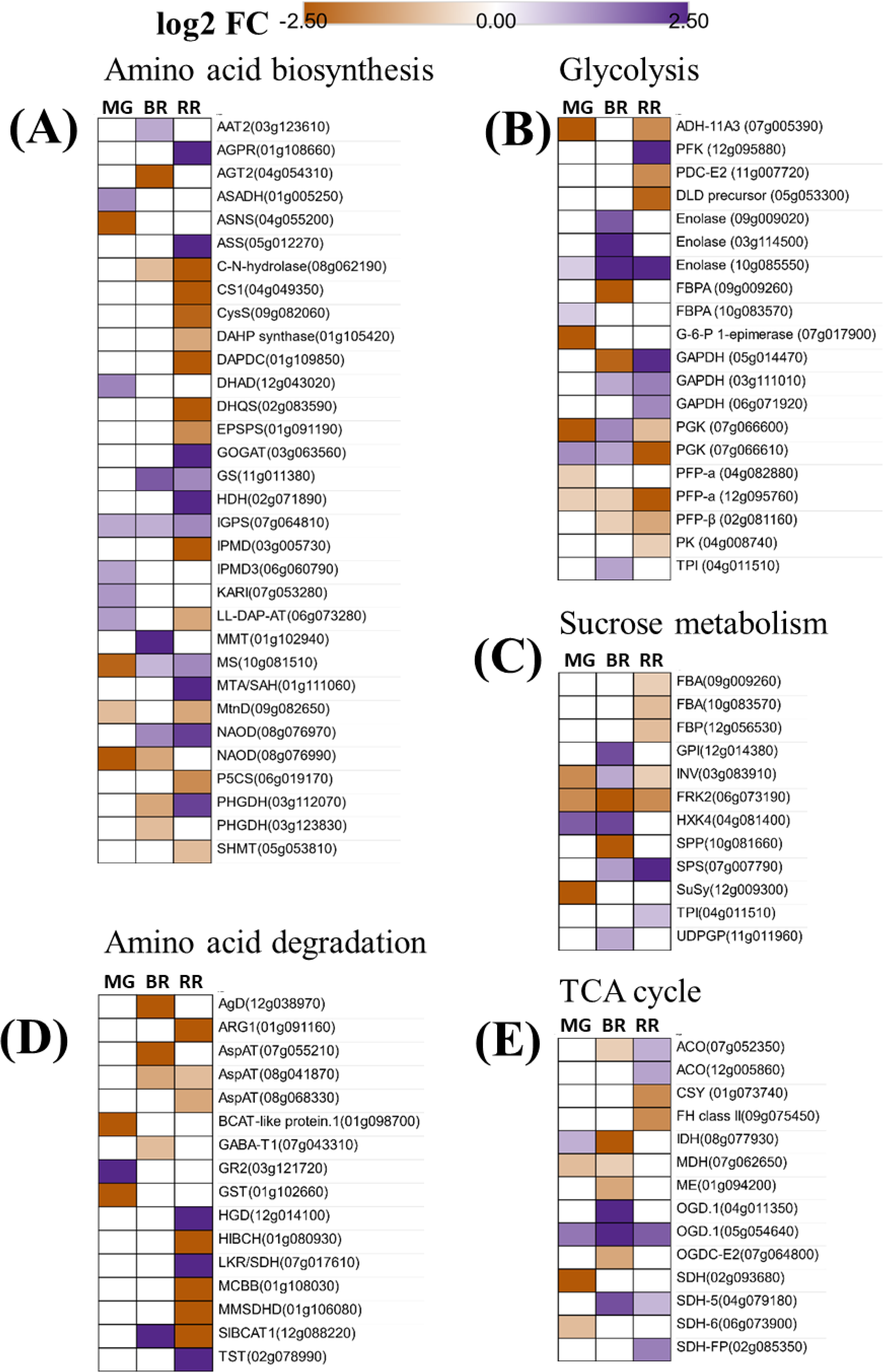
Heat maps representation of differentially expressed proteins affecting metabolic pathways in PKM-1 during fruit ripening. The altered proteins are shown for **(A)** amino acid biosynthesis, **(B)** glycolysis, **(C)** sucrose metabolism, **(D)** amino acid degradation, and **(E)** TCA cycle. Only significantly different proteins (log2 fold ≥±0.58, P ≤ 0.05) are shown in heat maps. The SOLYC prefix before each protein number is removed for convenience. Data are means ± SE (n = 3), P ≤ 0.05. See Dataset 3 for detailed proteome data and the abbreviations of the proteins.

**Fig. S7.**
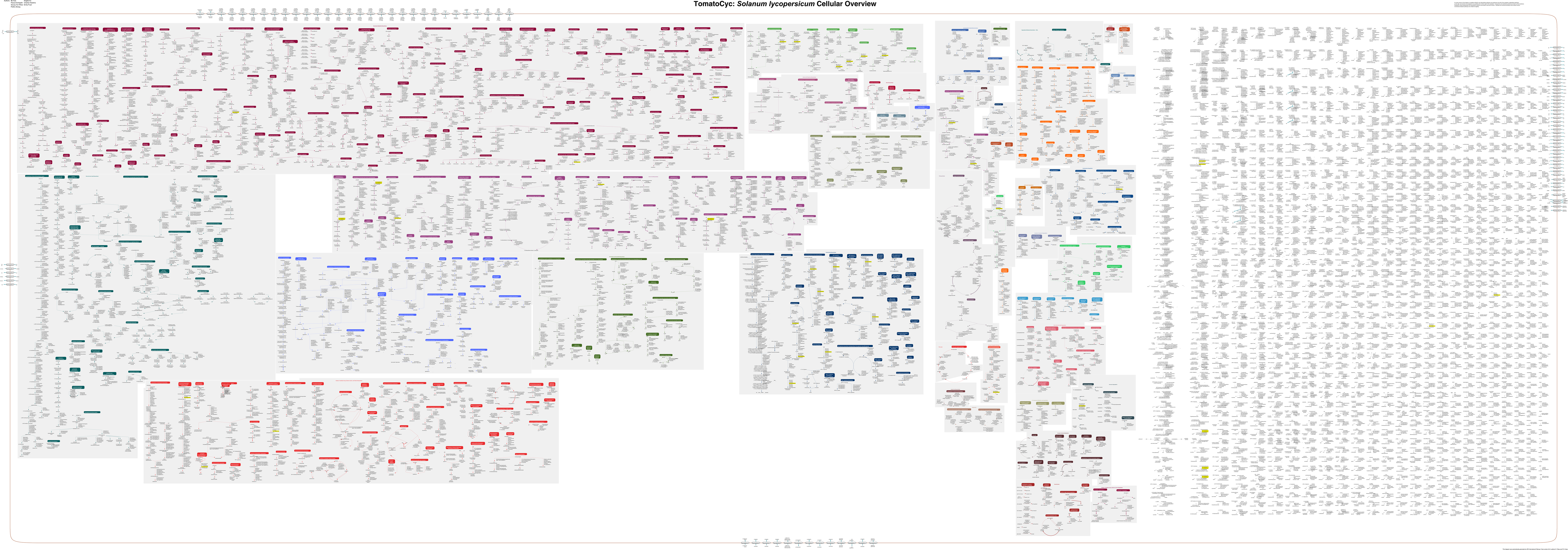
The tomato metabolic pathway marked with deleterious genes predicted by SIFT in PKM-1. The genes are highlighted in yellow color on the pathway. (For details, see **Dataset 4).**

**Table S1.**
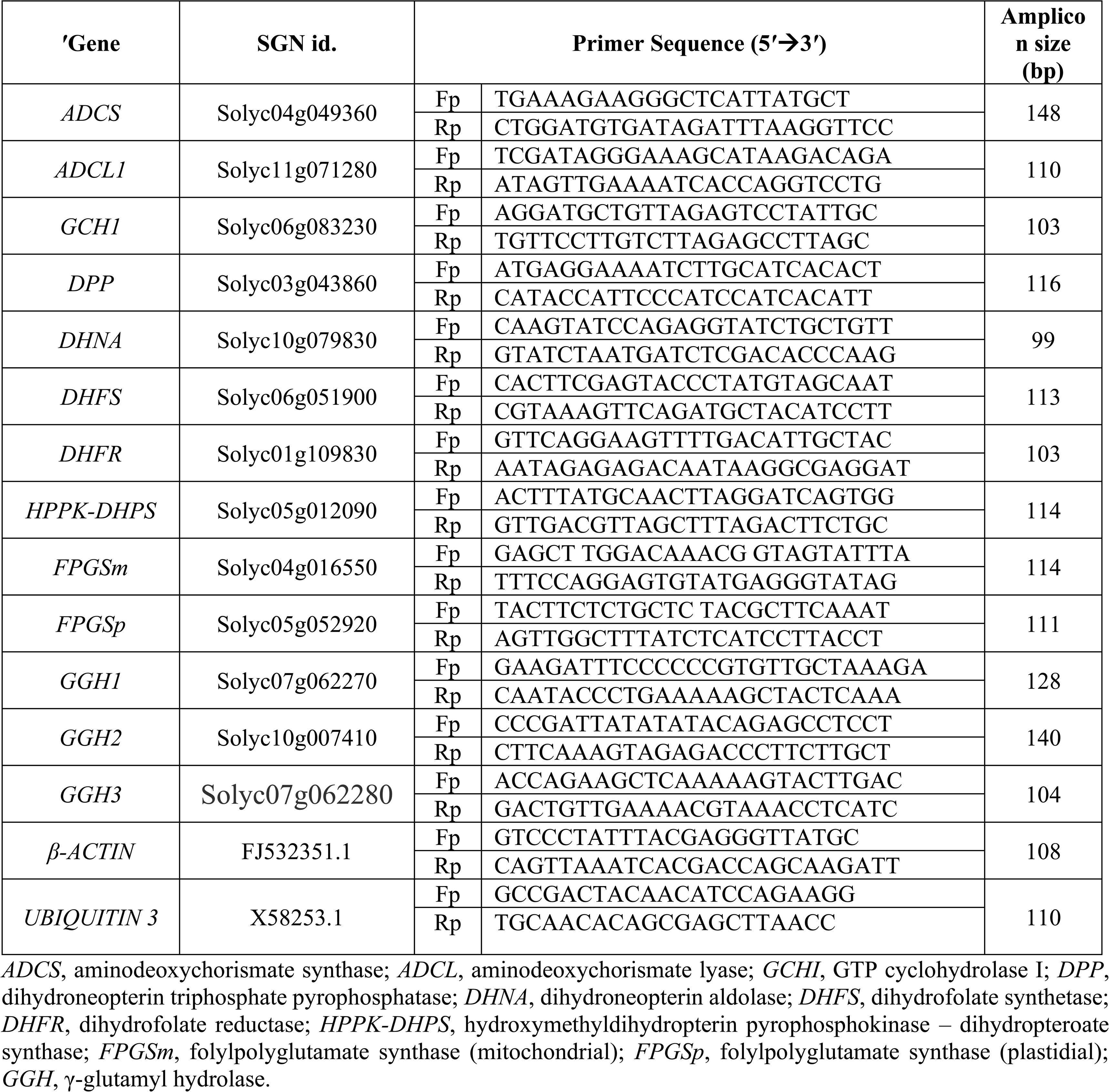
List of primers used for qRT-PCR analysis for folate biosynthesis pathway in AV and PKM-1 fruit. The primers were designed using SOL ITAG 2.3.

**Table S2.**
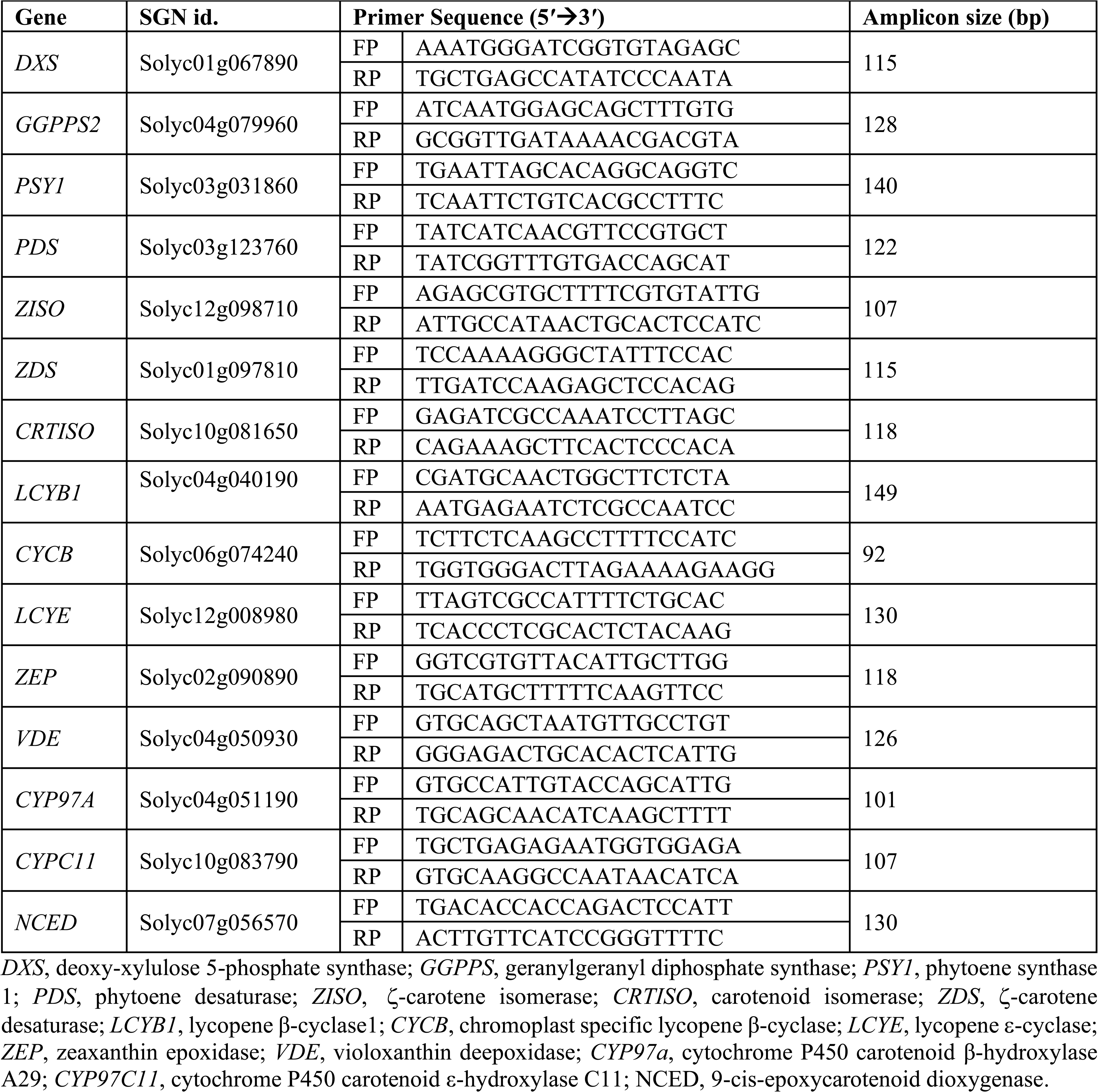
List of primers used for qRT-PCR analysis for carotenoid biosynthesis pathway in AV and PKM-1 fruit. The primers were designed using SOL ITAG 2.3

**Table S3.**
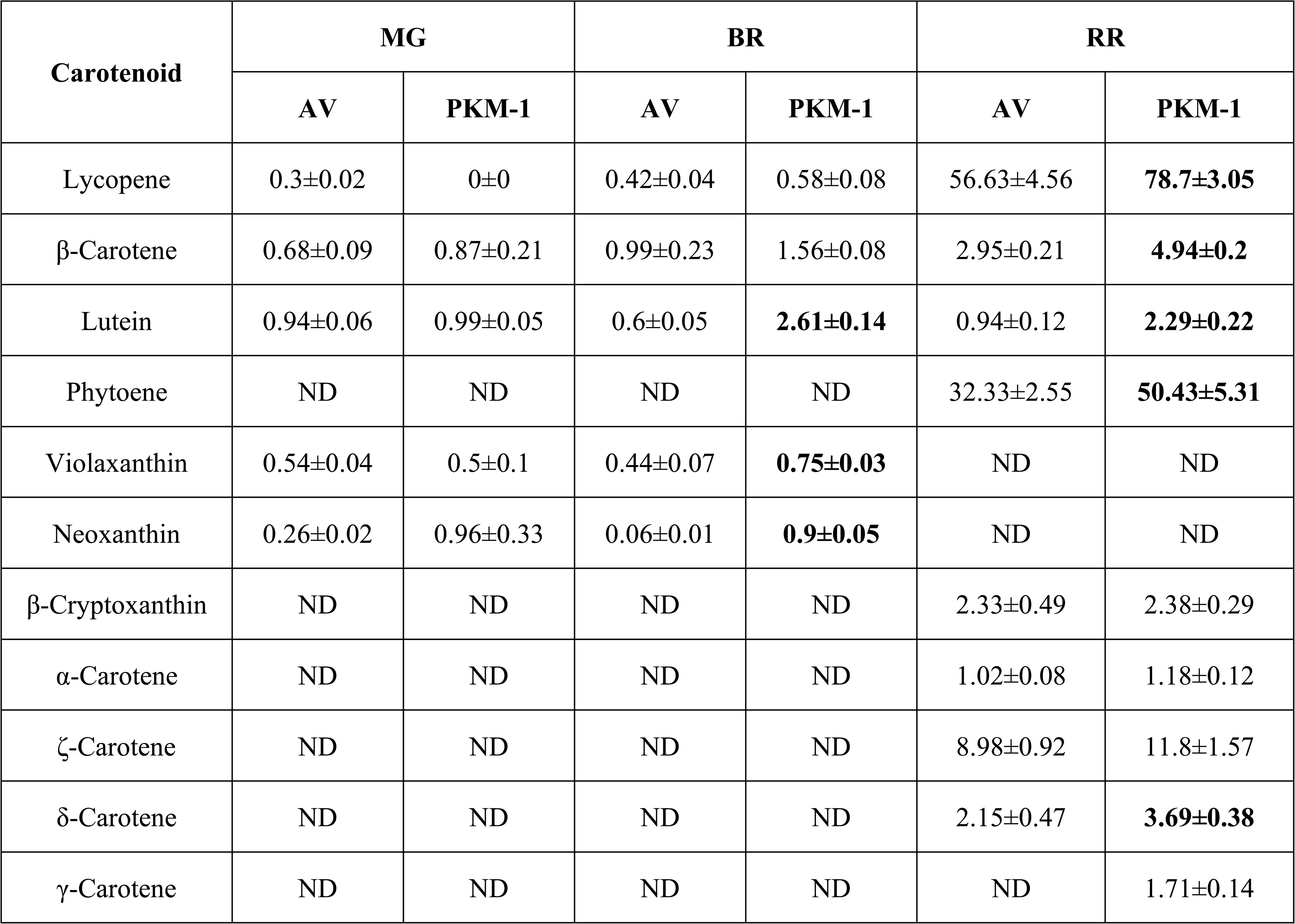
Carotenoids profiles of AV and PKM-1 at MG, BR, and RR fruits. Values in bold show the significant difference (P ≤ 0.05) with respect to AV. Data are mean ± SE (n =4) and expressed as µg/gm FW. The ND (Not detected) refers to the stages where the carotenoid analysis was carried out, but respective carotenoid was not detected, perhaps due to being below the limit of the detection.

**Table S4.**
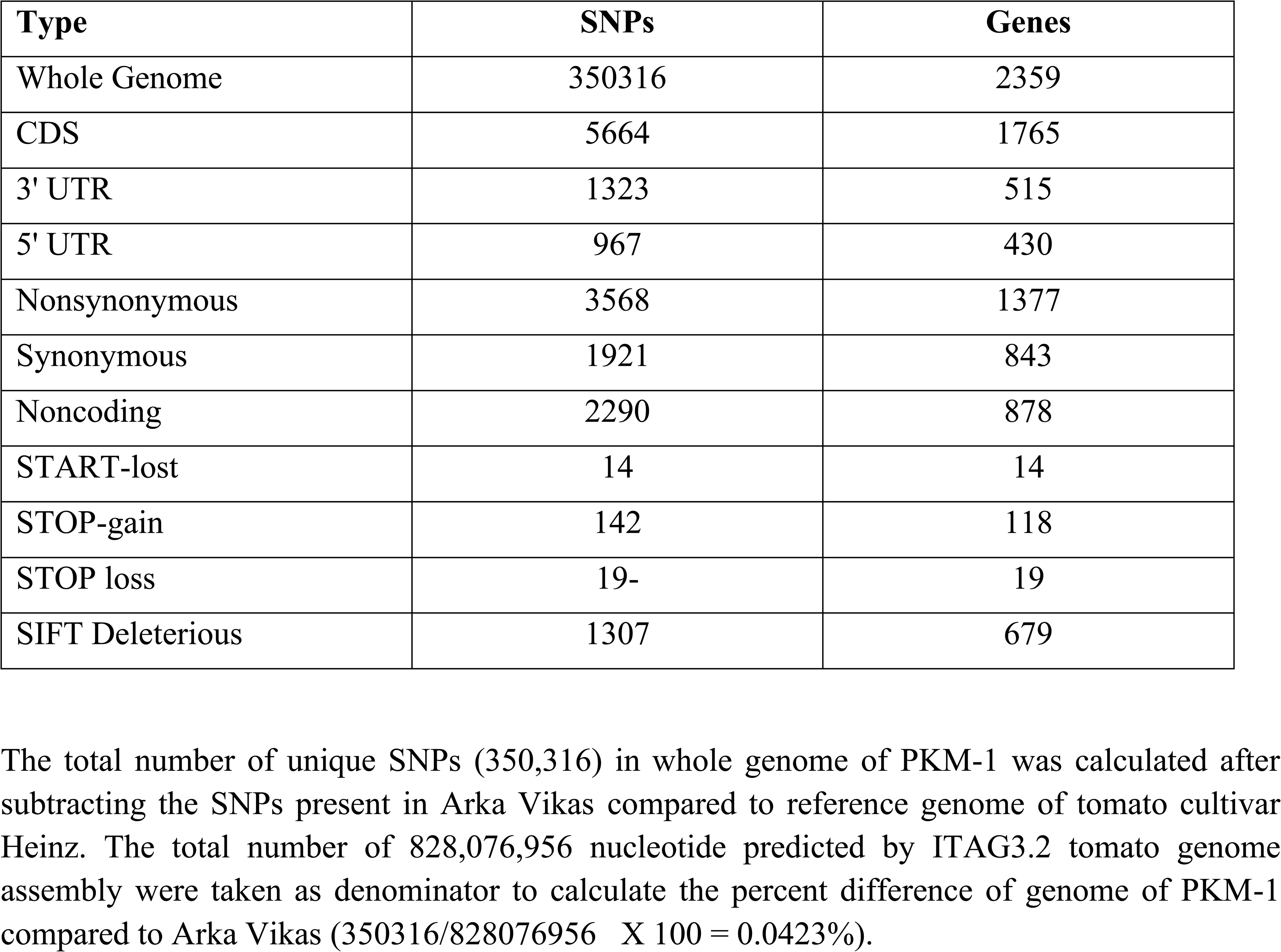
The relative proportions of different functional categories of mutations in genes and in noncoding regions including promoters and introns in PKM-1.

## References

1. Akhtar TA, Orsomando G, Mehrshahi P, Lara-Núñez A, Bennett MJ, Gregory III JF, Hanson AD, (2010) A central role for gamma-glutamyl hydrolases in plant folate homeostasis. Plant Journal, 64, 256–266.

2. Bali S, Robinson BR, Sathuvalli V, Bamberg J, Goyer A, (2018) Single nucleotide polymorphism (SNP) markers associated with high folate content in wild potato species. PLoS One.;13(*2*), e0193415.

3. Basset GJ, Ravanel S, Quinlivan EP, White R, Giovannoni JJ, Rébeillé F, Nichols BP, Shinozaki K, Seki M, Gregory III JF, Hanson AD, (2004) Folate synthesis in plants: the last step of the p-aminobenzoate branch is catalyzed by a plastidial aminodeoxychorismate lyase. Plant Journal, 40, 453–461.

4. Blancquaert D, De Steur H, Gellynck X, Van Der Straeten D, (2014) Present and future of folate biofortification of crop plants. Journal of Experimental Botany. 65, 895–906.

5. Blancquaert D, Storozhenko S, Van Daele J, Stove C, Visser RG, Lambert W, Van Der Straeten D, (2013) Enhancing pterin and para-aminobenzoate content is not sufficient to successfully biofortify potato tubers and Arabidopsis thaliana plants with folate. Journal of Experimental Botany. 64, 3899–909.

6. Bodanapu R, Gupta SK, Basha PO, Sakthivel K, Sreelakshmi Y, Sharma R, (2016) Nitric oxide overproduction in tomato *shr* mutant shifts metabolic profiles and suppresses fruit growth and ripening. Frontiers in Plant Science, 7, p.1714.

7. Chaudhary P, Sharma A, Singh B, Nagpal AK, (2018) Bioactivities of phytochemicals present in tomato. Journal of Food Science and Technology, 55, 2833–2849.

8. de La Garza RID, Gregory JF, Hanson AD, (2007) Folate biofortification of tomato fruit. Proceedings of the National Academy of Sciences, 104, 4218–4222.

9. De Lepeleire J, Strobbe S, Verstraete J, Blancquaert D, Ambach L, Visser RG, Stove C, Van Der Straeten D, (2018) Folate biofortification of potato by tuber-specific expression of four folate biosynthesis genes. Molecular Plant, 11, 175–188.

10. Deruère J, Römer S, d’Harlingue A, Backhaus RA, Kuntz M, Camara B, (1994) Fibril assembly and carotenoid overaccumulation in chromoplasts: a model for supramolecular lipoprotein structures. Plant Cell. 6,11 9–33.

11. Fàbregas N, Fernie AR, (2021) The interface of central metabolism with hormone signaling in plants. Current Biology. 31, R1535–48.

12. Fraser PD, Enfissi EM, Halket JM, Truesdale MR, Yu D, Gerrish C, Bramley PM, (2007) Manipulation of phytoene levels in tomato fruit: effects on isoprenoids, plastids, and intermediary metabolism. Plant Cell, 19, 3194–211

13. Gorelova V, Ambach L, Rébeillé F, Stove C, Van Der Straeten D, (2017a) Folates in plants: research advances and progress in crop biofortification. Frontiers In Chemistry, 5, 21.

14. Gorelova V, De Lepeleire J, Van Daele J, Pluim D, Meï C, Cuypers A, Leroux O, Rébeillé F, Schellens JH, Blancquaert D, Stove CP, (2017b) Dihydrofolate reductase/thymidylate synthase fine-tunes the folate status and controls redox homeostasis in plants. Plant Cell, 29, 2831–2853.

15. Guo W, Lian T, Wang B, Guan J, Yuan D, Wang H, Safiul Azam FM. Wan X, Wang W, Liang Q, Wang H, (2019) Genetic mapping of folate QTLs using a segregated population in maize. Journal of Integrative Plant Biology, 61, 675–690.

16. Gupta P, Dholaniya PS, Devulapalli S, Tawari NR, Sreelakshmi Y, Sharma R, (2020) Reanalysis of Genome Sequences of tomato accessions and its wild relatives: Development of tomato genomic variation (TGV) database integrating SNPs and INDELs polymorphisms. Bioinformatics 36, 4984–4990.

17. Gupta P, Dholaniya PS, Princy K, Madhavan AS, Sreelakshmi Y, Sharma R (2022) Whole-Genome Profiling of Ethyl Methanesulfonate Mutagenesis in Tomato. BioRixv https://doi.org/10.1101/2022.04.19.488728

18. Gupta P, Reddaiah B, Salava H, Upadhaya P, Tyagi P, Sarma S, Datta S, Malhotra B, Thomas T, Sunkum A, Devulapalli S, Till BJ, Sreelakshmi Y, Sharma R (2017) NGS-based identification of induced mutations in a doubly mutagenized tomato (*Solanum lycopersicum*) population. Plant Journal 92: 495–508

19. Gupta P, Sreelakshmi Y, Sharma R, (2015) A rapid and sensitive method for determination of carotenoids in plant tissues by high performance liquid chromatography. Plant Methods, 11, 1–12.

20. Gupta SK, Sharma S, Santisree P, Kilambi HV, Appenroth K, Sreelakshmi Y, Sharma R, (2014) Complex and shifting interactions of phytochromes regulate fruit development in tomato. Plant, Cell and Environment, 37, 1688–1702.

21. Hanson AD, Roje S, (2001) One-carbon metabolism in higher plants. Annual Review of Plant Biology, 52, 119–137.

22. Houben M, Van de Poel B, (2019) 1-Aminocyclopropane-1-Carboxylic Acid Oxidase (ACO): The Enzyme That Makes the Plant Hormone Ethylene Frontiers in Plant Science, 10, 695

23. Iniesta MD, Perez-Conesa D, Garcia-Alonso J, Ros G, Periago MJ, (2009) Folate content in tomato (Lycopersicon esculentum). Influence of cultivar, ripeness, year of harvest, and pasteurization and storage temperatures. Journal of Agricultural and Food Chemistry, 57, 4739–4745.

24. Jiang L, Liu Y, Sun H, Han Y, Li J, Li C, Guo W, Meng H, Li S, Fan Y, Zhang C, (2013) The mitochondrial folylpolyglutamate synthetase gene is required for nitrogen utilization during early seedling development in Arabidopsis. Plant Physiology, 161, 971–989.

25. Jiang L, Strobbe S, Van Der Straeten D, Zhang C, (2021) Regulation of plant vitamin metabolism: backbone of biofortification for the alleviation of hidden hunger. Molecular Plant. 14:40–60

26. Kilambi HV, Dindu A, Sharma K, Nizampatnam NR, Gupta N, Thazath NP, Dhanya AJ, Tyagi K, Sharma S, Sharma R, Sreelakshmi YS (2021) The new kid on the block: A dominant-negative mutation of phototropin1 enhances carotenoid content in tomato fruits. Plant Journal 106, 844–6

27. Kilambi HV, Kumar R, Sharma R, Sreelakshmi Y, (2013) Chromoplast-specific carotenoid-associated protein appears to be important for enhanced accumulation of carotenoids in hp1 tomato fruits. Plant Physiology, 161, 2085–2101.

28. Kilambi HV, Manda K, Sanivarapu H, Maurya VK, Sharma R, Sreelakshmi Y (2016) Shotgun Proteomics of Tomato Fruits: Evaluation, Optimization and Validation of Sample Preparation Methods and Mass Spectrometric Parameters. Frontiers in Plant Science. 7, 969.

29. Koike, S., Matsukura, C., Takayama, M., Asamizu, E., Ezura, H, (2013) Suppression of γ- aminobutyric acid (GABA) transaminases induces prominent GABA accumulation, dwarfism and infertility in the tomato (*Solanum lycopersicum* L.) Plant Cell Physiol. 54, 793–807.

30. Lanahan MB, Yen HC, Giovannoni JJ, Klee HJ, (1994) The never ripe mutation blocks ethylene perception in tomato. The Plant Cell. 6:521–30.

31. Li H, Durbin R. (2009) Fast and accurate short read alignment with Burrows–Wheeler transform. Bioinformatics 25,1754–60.

32. Li W, Liang Q, Mishra RC, Sanchez-Muñoz R, Wang H Chen, X, Van Der Straeten D, Zhang C, Xiao Y, (2021) The 5-formyl-tetrahydrofolate proteome links folates with C/N metabolism and reveals feedback regulation of folate biosynthesis. The Plant Cell. 33, 3367–85

33. Lian T, Guo W, Chen M, Li J, Liang Q, Liu F, Meng H, Xu B, Chen J, Zhang C, Jiang L. (2015) Genome-wide identification and transcriptional analysis of folate metabolism- related genes in maize kernels. BMC Plant Biology. 15, 204.

34. Liu L, Shao Z, Zhang M, Wang Q, (2015) Regulation of carotenoid metabolism in tomato. Molecular Plant 8, 28–39.

35. Martínez-Reyes I., Chandel NS, (2020) Mitochondrial TCA cycle metabolites control physiology and disease. Nature Communications, 11, 1–11.

36. Martín-Tornero E, Gómez DG, Durán-Merás I, Espinosa-Mansilla A, (2016) Development of an HPLC-MS method for the determination of natural pteridines in tomato samples. Analytical Methods, 8, 6404–6414.

37. Mehrshahi P, Gonzalez-Jorge S, Akhtar TA, Ward JL, Santoyo-Castelazo A, Marcus SE, Lara-Núñez A, Ravanel S, Hawkins ND, Beale MH, Barrett DA, (2010) Functional analysis of folate polyglutamylation and its essential role in plant metabolism and development. Plant Journal, 64, 267–279.

38. Meyer RS, Purugganan MD (2013) Evolution of crop species: genetics of domestication and diversification. Nature Review Genetics, 14, 840–852,

39. Mohan V, Gupta S, Thomas S, Mickey H, Charakana C, Chauhan VS, Sharma K, Kumar R, Tyagi K, Sarma S, Gupta SK, Kilambi HV, Nongmaithem S, Kumari A, Gupta P, Sreelakshmi Y, Sharma R (2016) Tomato fruits show wide phenomic diversity but fruit developmental genes show low genomic diversity. PLoS ONE 11, e0152907.

40. Navarrete O, Van Daele J, Stove C, Lambert W, Van Der Straeten D, Storozhenko S, (2012) A folate independent role for cytosolic HPPK/DHPS upon stress in Arabidopsis thaliana. Phytochemistry, 73, 23–33.

41. Piironen V, Edelmann M, Kariluoto S, Bedo Z, (2008) Folate in wheat genotypes in the HEALTHGRAIN diversity screen. Journal of Agricultural and Food Chemistry, 56, 9726–9731.

42. Pourcq KD, (2014) New insights into carotenoid accumulation in tomato fruit. PhD thesis. University of Barcelona

43. Shane B (1989) Folylpolyglutamate synthesis and role in the regulation of one-carbon metabolism. Vitamins & Hormones. 45, 263–335

44. Sharma K, Gupta S, Sarma S, Rai M, Sreelakshmi Y, Sharma R, (2021) Mutations in Tomato 1-Aminocyclopropane Carboxylic Acid Synthase2 Uncover Its Role in Development beside System II Responses. Plant Journal 106, 95–112.

45. Shohag MJI, Wei YY, Yu N, Zhang J, Wang K, Patring J, He ZL, Yang XE, (2011) Natural variation of folate content and composition in spinach (Spinacia oleracea) germplasm. Journal of Agricultural and Food Chemistry, 59, 12520–12526.

46. Simkin AJ, Gaffé J, Alcaraz JP, Carde JP, Bramley PM, Fraser PD, Kuntz M, (2007) Fibrillin influence on plastid ultrastructure and pigment content in tomato fruit. Phytochemistry, 68, 1545–1556.

47. Skirycz A, Caldana C, Fernie AR, (2022) Regulation of Plant Primary Metabolism–How Results From Novel Technologies Are Extending Our Understanding From Classical Targeted Approaches. Critical Reviews in Plant Sciences. 23:1–20.

48. Storozhenko S, De Brouwer V, Volckaert M, Navarrete O, Blancquaert D, Zhang GF, Lambert W, Van Der Straeten D, (2007) Folate fortification of rice by metabolic engineering. Nature Biotechnology, 25, 1277–1279.

49. Stover P, Schirch V, (1993) The metabolic role of leucovorin. Trends in Biochemical Sciences, 18, 102–106.

50. Strålsjö L, Alklint C, Olsson ME, Sjöholm I, (2003) Total folate content and retention in rosehips (Rosa ssp.) after drying. Journal of Agricultural and Food Chemistry. 51, 4291–5.

51. Theologis A, Oeller PW, Wong LM, Rottmann WH, Gantz DM, (1993) Use of a tomato mutant constructed with reverse genetics to study fruit ripening, a complex developmental process. Developmental Genetics. 14, 282–95.

52. Tyagi K, Sunkum A, Rai M, Yadav A, Sircar S, Sreelakshmi Y, Sharma R, (2022) Seeing the unseen: A trifoliate (MYB117) mutant allele fortifies folate and carotenoids in tomato fruits. Plant Journal https://doi.org/10.1111/tpj.15925

53. Tyagi K, Upadhyaya P, Sarma S, Tamboli V, Sreelakshmi Y, Sharma R, (2015) High performance liquid chromatography coupled to mass spectrometry for profiling and quantitative analysis of folate monoglutamates in tomato. Food Chemistry, 179, 76–84.

54. Upadhyaya P, Tyagi K, Sarma S, Tamboli V, Sreelakshmi Y, Sharma R, (2017) Natural variation in folate levels among tomato (Solanum lycopersicum) accessions. Food Chemistry, 217, 610–619.

55. Verwoerd TC, Dekker BM, Hoekema A, (1989) A small-scale procedure for the rapid isolation of plant RNAs. Nucleic Acids Research, 17, 2362.

56. Vishnevetsky M, Ovadis M, Vainstein A, (1999) Carotenoid sequestration in plants: the role of carotenoid-associated proteins. Trends in Plant Science. 4, 232–5.

57. Vizcaíno JA, Deutsch EW, Wang R, Csordas A, Reisinger F, Ríos D, Dianes JA, Sun Z, Farrah T, Bandeira N, Binz PA, (2014) ProteomeXchange provides globally coordinated proteomics data submission and dissemination. Nature Biotechnology. 32, 223–6.

58. Waller JC, Akhtar TA, Lara-Núñez A, Gregory III JF, McQuinn RP, Giovannoni JJ, Hanson AD, (2010) Developmental and feedforward control of the expression of folate biosynthesis genes in tomato fruit. Molecular Plant. 3, 66–77.

59. Waltz E, (2022) GABA-enriched tomato is first CRISPR-edited food to enter market. Nature Biotechnology, 40, 9–11.

60. Wang L, Kong D., Lv, Q., Niu, G., Han, T., Zhao, X., Meng, S., Cheng, Q., Guo, S., Du, J. and Wu, Z., (2017) Tetrahydrofolate modulates floral transition through epigenetic silencing. Plant Physiology, 174, 1274–1284.

61. Wei DO, Cheng ZJ, Xu JL, Zheng TQ, Wang XL, Zhang HZ, Jie WA, Wan JM (2014) Identification of QTLs underlying folate content in milled rice. Journal of Integrative Agriculture. 13, 1827–34.

62. Zanor MI, Osorio S, Nunes-Nesi A, Carrari F, Lohse M, Usadel B, Kuhn C, Bleiss W, Giavalisco P, Willmitzer L, Sulpice R, (2009) RNA interference of LIN5 in tomato confirms its role in controlling Brix content, uncovers the influence of sugars on the levels of fruit hormones, and demonstrates the importance of sucrose cleavage for normal fruit development and fertility. Plant Physiology. 150, 1204–18.

